# The dissociable effects of reward on sequential motor behaviour

**DOI:** 10.1101/2021.09.17.460761

**Authors:** Sebastian Sporn, Xiuli Chen, Joseph M Galea

**Affiliations:** School of Psychology and Centre for Human Brain Health, University of Birmingham, UK; Department of Clinical and Movement Neuroscience, Queens Square Institute of Neurology, UCL, UK

**Author notes:** <u>Correspondence:</u> Sebastian Sporn.

## Abstract

Reward has consistently been shown to enhance motor performance however its beneficial effects appear to be largely unspecific. While reward has been shown to invigorate performance, it also enhances learning and/or retention. Therefore, a mechanistic account of the effects of reward on motor behaviour is lacking. Here we tested the hypothesis that these distinct reward-based improvements are driven by dissociable reward types: explicit reward (i.e. money) and performance feedback (i.e. points). Experiment 1 showed that explicit reward instantaneously improved movement times (MT) using a novel sequential reaching task. In contrast, performance-based feedback led to learning-related improvements. Importantly, pairing both maximised MT performance gains and accelerated movement fusion. Fusion describes an optimisation process during which neighbouring sequential movements blend together to form singular actions. Results from experiment 2 served as a replication and showed that fusion led to enhanced performance speed whilst also improving movement efficiency through increased smoothness. Finally, experiment 3 showed that these improvements in performance persist for 24 hours even without reward availability. This highlights the dissociable impact of explicit reward and performance feedback, with their combination maximising performance gains and leading to stable improvements in the speed and efficiency of sequential actions.

## Introduction

Research into the effects of reward on motor behaviour has consistently shown that reward enhances performance^1–11^. Consequently reward, as a tool to shape motor behaviour, has gained much scientific interest particularly with regards to its strategic and beneficial use in rehabilitation. However, the beneficial effects of reward on behaviour appear to be largely unspecific. While studies using saccadic or discrete reaching movements have consistently found that reward instantaneously improved the speed-accuracy function (i.e. transient improvement within a single trial)^1,3–5,11–14^, research employing force and button-press tasks showed reward-related improvements in learning and/or retention (i.e. improvements across trials)^8–10,15–17^. Therefore, a mechanistic account for how reward enhances motor performance is lacking, which restricts the potential of a targeted use of reward in clinical settings. Crucially, such a mechanistic account will have to be able to account for both the invigoration and training-dependent learning effects associated with reward.

An interesting possibility is that these distinct reward-based improvements are driven by dissociable reward types. Reward is most commonly provided through either explicit reward (i.e., money), performance feedback (i.e., points) or a combination of both^4,8–10,15^. Performance feedback represents a reinforcement-based teaching signal (i.e., reward-prediction error)^18,19^ which provides information on how well a motor task has been completed (knowledge of performance) and has been shown to enhance other forms of motor learning^11,20–24^. In contrast, explicit reward such as a monetary incentive increases the motivation to perform optimally without necessarily providing a performance-based learning signal^25^. Recent research decoupled explicit reward from performance feedback and found that while performance feedback alone was not sufficient to induce skill leaning in a pinch force reproduction task, combining it with explicit reward was^15^. However, the effect of explicit reward alone on motor behaviour was not accounted for^15^. Therefore, to dissociate the effects of both on motor behaviour it is crucial to systematically assess them in isolation and in combination.

To this end, we designed a novel complex sequential motor task in which participants were asked to complete a continuous sequence of 8 reaching movements. Participants received a combination of explicit reward (money) and performance feedback (points), which allowed us to systematically evaluate how they influence performance during a complex sequential reaching task. We hypothesised that explicit reward, which has been shown to increase motivation, will instantaneously enhance performance (i.e., during early training). In contrast, accurate performance feedback will lead to learning-related improvements across training. Importantly, in line with recent findings, we hypothesised that combining explicit reward with accurate performance feedback will maximise performance gains^15^.

Experiment 1 confirmed that explicit reward and performance feedback have dissociable effects on motor behaviour. Specifically, participants who received explicit reward, irrespective of the availability and quality of performance-based feedback, instantly reduced movement times (MT) during early training. Additionally, accurate performance feedback led to training-related improvements in MT irrespective of reward availability. Crucially, combining explicit reward with accurate performance feedback resulted in both an instant reduction and a learning-related improvement in MT which maximised performance gains. Further analysis revealed that these performance gains were associated to movement fusion. Fusion describes an optimisation process during which individual motor elements are blended into a combined singular action^26–28^. Therefore, movement fusion represents an effective strategy to achieve quicker MTs by producing fast reaching movements while simultaneously reducing dwell times when transitioning between reaches. Experiment 2 provided a replication in which the combination of explicit reward and accurate performance feedback improved MTs across 2 days of training and led to a substantial increase in movement fusion. Critically, movement fusion was associated with increases in movement smoothness which also reflected the predictions of a model that optimised jerk across the sequential movement^29,30^. These results suggest that performance became more energetically-efficient^31,32^ which may explain the results from experiment 3 where improvements in performance persisted for 24 hours even when reward was no longer available.

## Results

To assess the influence of explicit reward and performance-based feedback on complex, sequential movements we developed a novel reaching task in which participants made 8 sequential reaching movements to designated targets (1 trial) using a motion tracking device (Figure 1a, b). Prior to the start of the experiment, participants were trained on the sequence without a time constraint until reaching a learning criterion of 5 successful trials in a row. Importantly, missing a target resulted in an immediate abortion of the current trial, which participants then had to repeat. Therefore, any performance gains could only be attributed to improvements in the execution and not to memory of the sequence or decreases in accuracy. Participants then completed a baseline period (10 trials) during which they were encouraged to complete each trial ‘as fast and as accurately as possible’ (Figure 1c). Afterwards, participants completed 200 training trials. During training, participants were placed under different feedback regimes which differed with regards to both the availability of explicit reward (money) and performance-feedback (Figure 1d). This allowed us to systematically evaluate how they influence performance during a complex sequential reaching task.

**Figure 1.**
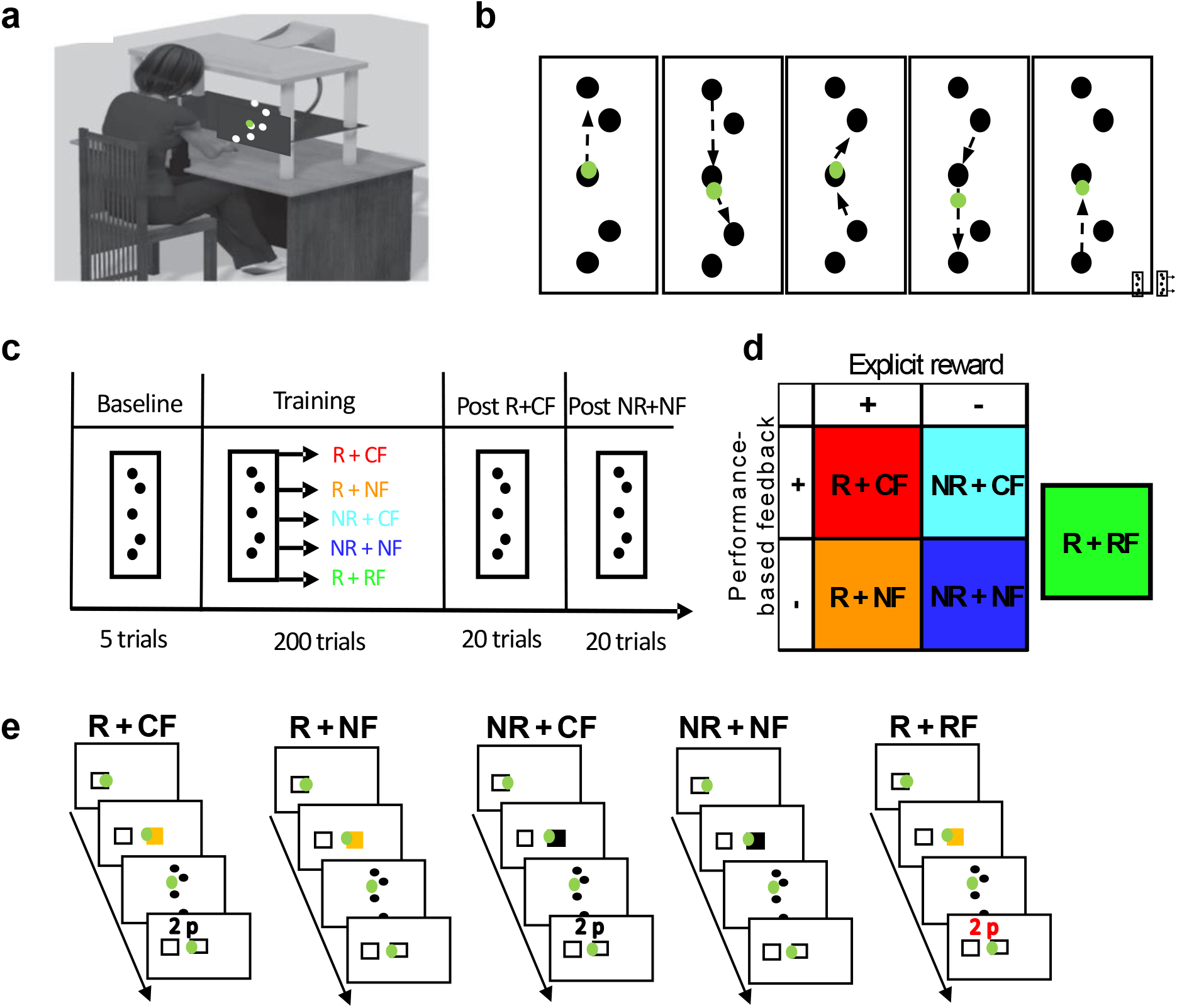
Experimental setup. **a)** Participants wore a motion-tracking device on the index finger and the unseen reaching movements were performed across a table whilst a green cursor matching the position of index finger was viewable on a screen. **b)** 8 movement sequential reaching task. Participants started from the centre target. **c)** Study design. Prior to the start of the experiment, participants were trained on the reaching sequence and were then asked to perform 10 baseline trials. Randomly allocated to one of five groups, participants completed 200 trials during training and an additional 20 trials in each post assessment; one with reward (post-R+CF) and one without (post-NR+NF) (counterbalanced across participants). **d)** Groups. The feedback regime differed with regards to the availability of both explicit reward (money) and performance-based feedback. Participants either received explicit reward (money) (R) or not (NR). Similarly, participants were either provided with correct (CF), no (NF) or random performance-based (RF) feedback. **e)** Explicit reward (money) trials were cued using a visual stimulus prior to the start of the trial. In contrast, in trials without explicit reward, participants were instructed to be as fast and accurate as possible. Feedback was provided once participants completed the trial and moved the cursor back to the start box.

Participants in Group_-R + CF_ (explicit reward + correct feedback) were able to earn money depending on their movement time (MT) and received performance-based feedback (the amount of money (0-5p) awarded in a given trial) after each trial. Explicit reward trials were cued with a yellow start box (Figure 1e), and the performance feedback was calculated using a closed-loop design comparing MT performance on a given trial with performance on the last 20 trials. This provided the participant with graded feedback (0-5p) of their MT performance relative to their recent performance (see Methods). In contrast, participants in Group_-R + NF_ (explicit reward + no feedback) only received explicit reward (yellow start box cue) but were not provided with performance feedback after each trial. Instead, participants received the accumulated monetary reward at the end of training. Similarly to Group_-R + CF_, participants in Group_-R + RF_ (explicit reward + random feedback) received both explicit reward and performance-based feedback. However, the performance feedback was random (see Methods) and thus did not correspond to participant’s actual performance. Finally, while participants in both Group_-NR + CF_ (no explicit reward + correct feedback) and Group_-NR + NF_ (no explicit reward + no feedback) did not receive any explicit reward during training, performance feedback was provided in Group_-NR + CF_. However, participants were told that this feedback was not related to monetary incentive (Figure 1e).

After training, both groups engaged in a rewarded (post-R+CF) and no rewarded (post-NR+NF) post-assessment (20 trials each). Therefore, all groups received both explicit reward and correct feedback during post-R+CF, while neither was available during post-NR+NF the post-assessment intended to compare performance between groups when under the same condition (Figure 1c).

### Monetary incentive led to an instantaneous decrease in MT while performance-based feedback was associated with learning-related MT improvements

MT reflected total movement duration from exiting the start box until reaching the last target. We found no difference at baseline (ANOVA; group, F = 1.35, p = 0.2603). To assess whether explicit reward and performance-based feedback have distinct effects on MT performance during training we computed 1000 bootstrap estimates of the data for each group. Each estimate represented a randomly generated data set (N=15) with replacement. A simple polynomial model (*f(x) = p1*x + p2*) was fit to the mean trial-by-trial training data (200 trials) for each bootstrap estimate. The 95% confidence intervals for each model parameter (*p1*: *gradient* & *p2: intercept*) were used to represent significant differences between groups^33^ (see Methods). Our results highlight that monetary incentive (explicit reward) instantaneously enhanced sequential reaching behaviour (Figure 2a, b). Specifically, groups that received explicit reward (Group_-R + CF_, MT = 4.3262, CI = [4.2162 4.4362]; Group_-R + NF_, MT = 4.6860, CI = [4.5616 4.8104]; Group_-R + RF_, MT = 4.3660, CI = [4.2567 4.4757]) exhibited lower MTs at the start of training than the NR groups (Group_-NR + NF_, MT = 5.138, CI = [5.0191 5.2570]; Group_-NR + CF_, MT = 5.3346, CI = [5.2389 5.4303]; Intercept, Figure 2b, c). In contrast, only groups that received accurate performance-based feedback (Group_-R + CF_, MT = -0.0072, CI = [-0.0078 -0.0066]; Group_-NR + CF_, MT = -0.0059, CI = [-0.0066 -0.0051]) showed greater learning-related decreases in MT across training, which suggests that feedback has to match performance to enhance learning (Group_-R + RF_, MT = -0.0028, CI = [-0.0034 -0.0022]; Slope, Figure 2b, d). Importantly, the combination of explicit reward (money) and accurate performance-based feedback maximised behavioural gains (Group_-R+CF_; Figure 2a, b). Note here that these improvements were not related to higher error rates which were of equal magnitude across groups and were consistently below an average of one error per trial across a group (Supplementary Figure 1).

**Figure 2.**
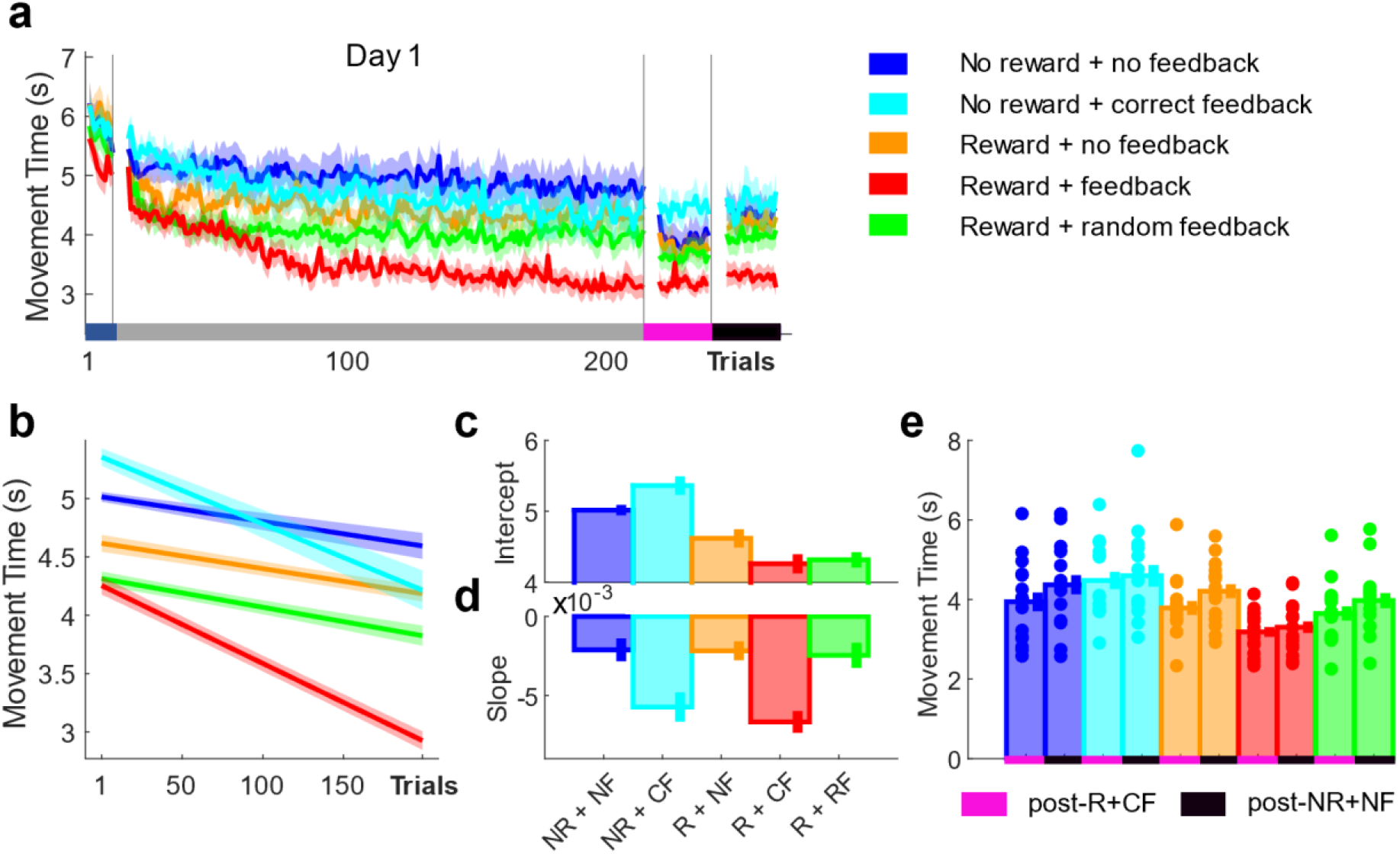
Explicit reward and performance-based feedback have distinct effects on MT. **a)** Trial-by-trial changes in MT averaged over participants for all groups. **b)** Averaged predicted model fits (simple polynomial model) to bootstrap estimates for each group including **e)** Intercept and **f)** Slope values (error bars represent 95% confidence intervals). **e)** Post assessment performance (post-R+CF vs post-NR+NF). Shaded regions/error bars represent SEM.

Across post assessments, we found a significant main effect for both timepoint (mixed-effect ANOVA; timepoint post-R (all 20 trials) vs post-NR (all 20 trials), F = 26.32, p < 0.0001; Figure 2e) and group (group, F = 4.56, p = 0.0025). Specifically, post hoc analysis revealed that Group_-R + CF_ was faster than both NR groups (Wilcoxon test; Group_-R + CF_ vs Group_-NR + NF_, Z = -1.3, p = 0.0011; Group_-R + CF_ vs Group_-NR + CF_, Z = 0.9, p = 0.0358). However, no further post hoc group comparisons yielded any significant results.

### Combined changes in maximum (vel_max_) and minimum (vel_min_) mediate improvements in MT

It has been shown that improvements in MT can be achieved via increases in maximum velocity (vel_max_) during simple discrete reaching movements^3,4^. To assess the effects of explicit reward and performance-based feedback on vel_max_ (see Methods, equation 1), we averaged vel_max_ across the 8 reaching movements (Figure 3a). We found no difference at baseline (ANOVA; group, F = 0.76, p = 0.5532) and an unclear result during early training (Intercept, Figure 3b, c).

**Figure 3.**
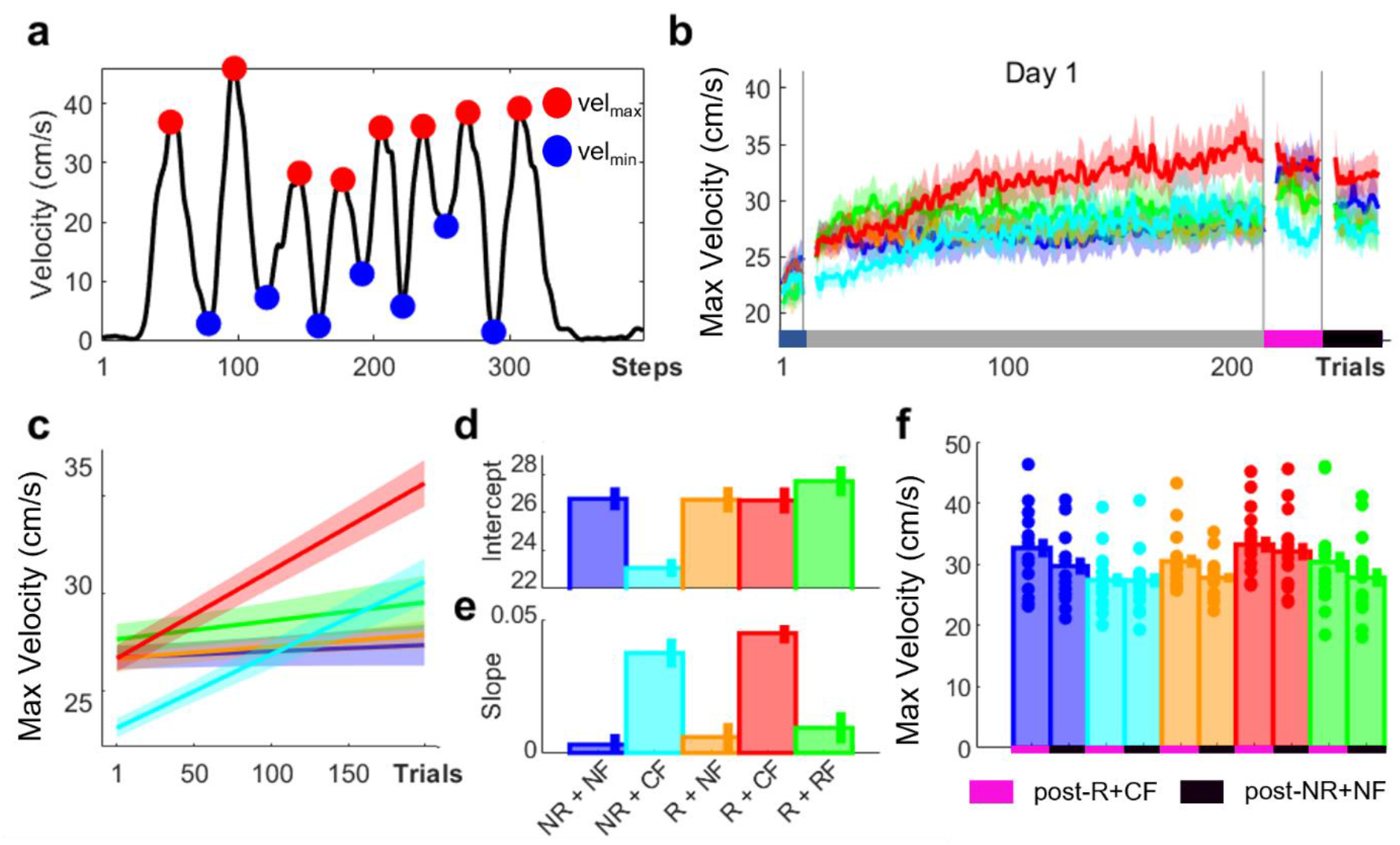
Accurate performance-based feedback led to training-dependent improvements in vel_max_. **a)** Illustration of vel_max_ and vel_min_. **b)** Trial-by-trial changes in vel_max_ averaged over participants for all groups. **c)** Averaged predicted model fits (simple polynomial model) to bootstrap estimates for each group including **e)** Intercept and **f)** Slope values (error bars represent 95% confidence intervals). **f)** Post assessment performance (post-R+CF vs post-NR+NF). Shaded regions/error bars represent SEM.

While all groups outperformed Group_-NR + CF_ (Group_-NR + CF_, vel_max_ = 23.3461, CI = [22.8807 23.8115]), Group_-NR + NF_ scored similar vel_max_ values to Group_-R + CF_ (Group_-NR + NF_, vel_max_ = 26.4338, CI = [25.8331 27.0345]; Group_-R + CF_, vel_max_ = 26.8560, CI = [26.1144 27.5976]). In contrast, accurate performance feedback was associated with a pronounced learning-related increase in vel_max_ (Group_-R + CF_, vel_max_ = 0.0438, CI = [0.0402 0.0474]; Group_-NR + CF_, vel_max_ = 0.0354, CI = [0.0305 0.0404]; Slope, Figure 3c, e). Importantly, and similarly to MT, the combination of explicit reward and accurate performance-based feedback maximised the gains observed in vel_max_ (Group_-R + CF_; Figure 2b, c). Across post assessments we found a significant main effect for timepoint (mixed-effect ANOVA; timepoint post-R vs post-NR, F = 31.25, p < 0.0001; Figure 2e) but not for group (group, F = 1.84, p = 0.1304).

In contrast to discrete motor behaviours, the task utilized for this study consisted of a sequence of reaching movements. Therefore, improvements in MT could additionally be driven by a reduction in dwell times when transitioning between reaching movements. To assess the effect of explicit reward and performance-based feedback on dwell times, we obtained vel_min_ and averaged them across the 7 reaching transitions (see Methods, equation 2; Figure 3a). Whilst no differences at baseline were observed (ANOVA; group, F = 1.26, p = 0.2923), explicit reward enhanced vel_min_ during early training (Group_-R + CF_, vel_min_ = 3.6630, CI = [3.4047 3.9212]; Group_-R + NF_, vel_min_ = 2.6834, CI = [2.3229 3.0439]; Group_-R + RF_, vel_min_ = 3.9792, CI = [3.5149 4.4436]; Intercept, Figure 4b, c). In contrast to vel_max_, Group_-NR+CF_ showed higher vel_min_ than Group_-NR+NF_ which were close to Group_-R+NF_ (Group_-NR + CF_, vel_min_ = 2.1124, CI = [1.7984 2.4263]). Additionally, performance feedback was associated with a learning-related increase in vel_min_ (Group_-R + CF_, vel_min_ = 0.0322, CI = [0.0290 0.0355]; Group_-NR + CF_, vel_min_ = 0.0160, CI = [0.0130 0.0190]; Slope, Figure 4b, d). Interestingly, random feedback in Group_-NR + RF_ led to similar learning slopes when compared to Group_-NR + CF_ (Group_-R + RF_, vel_min_ = 0.0126, CI = [0.0088 0.0165]), which may suggest that improvements in vel_min_ require feedback irrespective of whether it is accurate. Importantly, similarly to both MT and vel_max_, combining explicit reward with accurate performance feedback maximised the gains observed in vel_min_ (Group_-R + CF_; Figure 3a, c).

**Figure 4.**
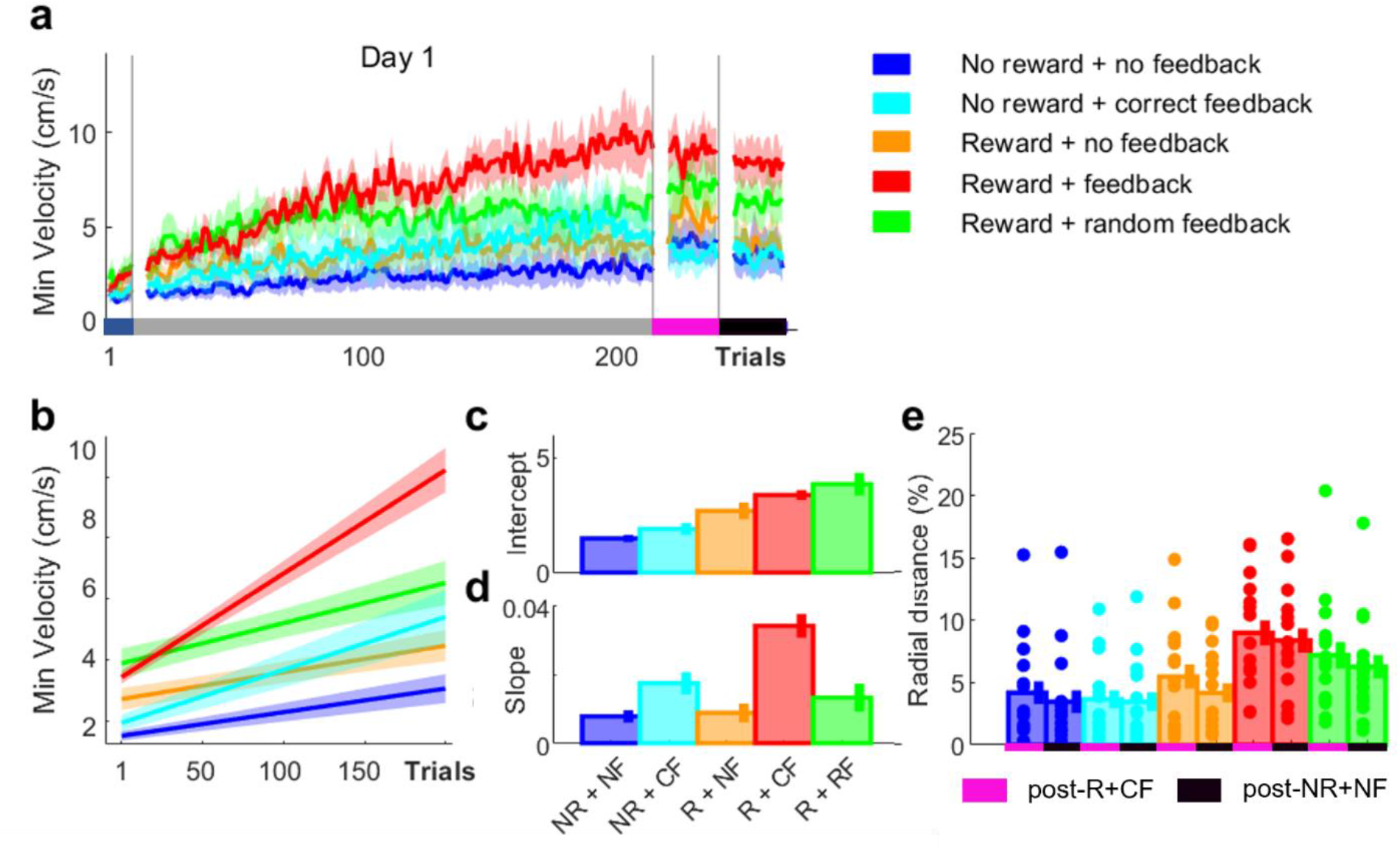
Accurate performance feedback led to training-dependent improvements in vel_min_. **a)** Trial-by-trial changes in vel_min_ averaged over participants for all groups. **b)** Averaged predicted model fits (simple polynomial model) to bootstrap estimates for each group including **e)** Intercept and **f)** Slope values (error bars represent 95% confidence intervals). **e)** Post assessment performance (post-R+CF vs post-NR+NF). Shaded regions/error bars represent SEM.

Across post assessments we found a significant main effect for both timepoint (mixed-effect ANOVA; timepoint post-R vs post-NR, F = 19.04, p < 0.0001; Figure 4e) and group (group, F = 4.41, p = 0.0003). Specifically, post hoc analysis revealed that Group_-R + CF_ had higher vel_min_ than both NR groups (Wilcoxon test; Group_-R + CF_ vs Group_-NR + NF_, Z = -0.81, p = 0.0108; Group_-R + CF_ vs Group_-NR + CF_, Z = 1.01, p = 0.0075). However, no further significant differences between groups were found. These results suggest that explicit reward (monetary incentive) and performance-based feedback have distinct effects on performance during a complex, sequential reaching task. Monetary incentive led to an instantaneous decrease in MTs while performance feedback was associated with a learning-dependent decrease in MT. This pattern was also observed in vel_max_ and vel_min_ which suggests that the combined changes in both underlie the MT results. Crucially, combining monetary incentive with accurate performance feedback maximised the behavioural gains observed.

### Movement fusion is associated with additional performance gains in MT

Thus, combining explicit reward with accurate performance-based feedback led to faster MTs by both increasing vel_max_ and vel_min_. One strategy to achieve faster reaching movements while simultaneously reducing dwell times when transitioning between reaches is movement fusion. Fusion describes the blending of individual motor elements into a combined action^26–28^. This is represented in the velocity profile by the stop period between two movements gradually disappearing and being replaced by a single velocity peak (Figure 5a). To measure movement fusion, we developed a fusion index (FI; Methods, Equation 3) that compared the mean vel_max_ of two sequential reaches with the vel_min_ around the via point (transition). The smaller the difference between these values, the greater fusion had occurred of these two movements as reflected by a FI value closer to 1 (Figure 5b). Considering that participants completed 7 transitions to complete a trial the maximum FI value was 7.

**Figure 5.**
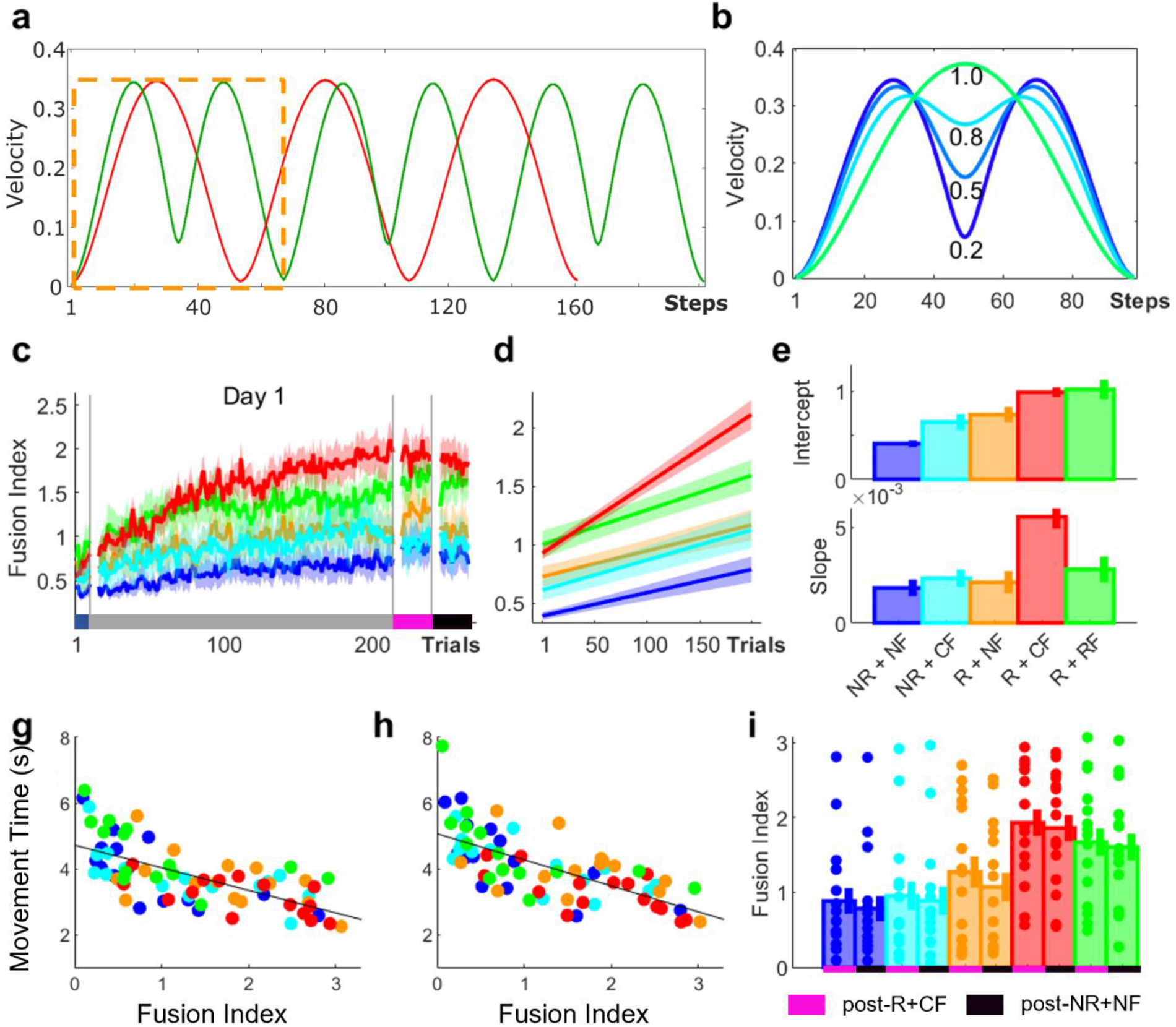
Movement fusion is strongly associated with improvements in MT. **a)** Illustration of a velocity profile corresponding to executing six reaching movements individually (green) and three fused reaches (red). Stop period, apparent when executing individual movements, disappears when executing a fully fused reaching movement (dashed orange). **b)** Illustration of fusion index (FI). **c)** Trial-by-trial changes in FI averaged over participants for all groups. **d)** Averaged predicted model fits (simple polynomial model) to bootstrap estimates for each group including **e)** Intercept and **f)** Slope values (error bars represent 95% confidence intervals). **g-h)** Scatterplots displaying the relationship between MT and FI levels during **g)** post-R+CF **h)** post-NR+NF with a linear line fitted across groups. **i)** Post assessment performance (post-R+CF vs post-NR+NF). Shaded regions/error bars represent SEM.

We found no difference at baseline (ANOVA; group, F = 1.36, p = 0.2555, Figure 5c). Explicit reward in combination with performance-based feedback (both correct and random) enhanced movement fusion during early training (Group_-R + CF_, FI = 0.9855, CI = [0.9280 1.0429]; Group_-R + RF_, FI = 1.0160, CI = [0.9075 1.1245]; Intercept, Figure 5d, e). In contrast, intercept differences between Group_-R + NF_ vs Group_-NR + CF_ were considerably smaller and closer to Group_-NR/NF_ (Group_-R + NF_, FI = 0.7225, CI = [0.6309 0.8141]; Group_-R + NF_, FI = 0.6419, CI = [0.5498 0.7341]; Group_-R + NF_, FI = 0.4034, CI = [0.3650 0.4419]). This suggests that early increases in movement fusion may depend on the availability of performance feedback in combination with reward. Importantly, in comparison to all other groups, only Group_-R + CF_ exhibited a pronounced learning-related increase in FI which was greater than in Group_-NR + CF_ and Group_-R + RF_ (Group_-R + CF_, FI = 0.0056, CI = [0.0050 0.0062]; Group_-NR + CF_, FI = 0.0022, CI = [0.0016 0.0028]; Group_-R + RF_, FI = 0.0028, CI = [0.0021 0.0035]). Furthermore, Group_-NR + CF_ and Group_-R + RF_ were not different from the NR groups (Group_-NR + CF_, FI = 0.0024, CI = [0.0019 0.0028]; Group_-NR + NF_, FI = 0.0018, CI = [0.0014 0.0022]; Slope, Figure 5d, f). Importantly, when correlating FI and MT during both post-R+CF (Figure 5g) and post-NR+NF (Figure 5h) we found that higher FI values were associated with MT while accounting for the factor group (post-R+CF; rho = -0.702, p < 0.0001; post-NR+NF; rho = -0.694, p < 0.0001). This highlights that faster MTs are related to increased fusion.

Across post assessments we found a significant main effect for both timepoint (mixed-effect ANOVA; timepoint post-R+CF vs post-NR+NF, F = 12.48, p < 0.0001; Figure 5i) and group (group, F = 4.85, p = 0.0017). Specifically, post hoc analysis revealed that Group-R + CF had higher FI values than both NR groups (Wilcoxon test; Group-R+CF vs Group-NR+NF, Z = -0.70, p = 0.0146; Group_-R+CF_ vs Group_-NR+CF_, Z = 1.06, p = 0.0050). However, no further significant differences were found.

### Explicit reward in combination with performance-based feedback led to significant improvements in performance across multiple days

To investigate whether these findings could be replicated and to further assess the kinematic underpinnings of movement fusion, we conducted a second experiment using the same task design. In experiment 2, only Group_-R + CF_ and Group_-NR + NF_ were included to contrast the most beneficial feedback regime (i.e., monetary incentive with accurate performance feedback with its logical opposite). Additionally, we added a further testing day (day 2) to investigate whether movement fusion can be further enhanced with additional training. During day 2, participants underwent the same experimental protocol as day 1 (Figure 1c).

We did not observe any differences during baseline for any measure (MT, Wilcoxon test; Z = -1.38, p = 0.17, Supplementary Figure 2a; vel_max_, Wilcoxon test; Z = 0.70, p = 0.4812, Supplementary Figure 2b; vel_min_, Wilcoxon test; Z = 1.44, p = 0.1516, Supplementary Figure 2c; FI, Wilcoxon test; Z = 1.31, p = 0.1908, Supplementary Figure 2d). Instead, we found that explicit reward in combination with performance-based feedback enhanced MT performance during early training on day 1 (Group_-R + CF_, MT = 3.8663, CI = [3.7725 3.9602], Group_-NR + NF_, MT = 4.9663, CI = [4.8428 5.0898], Intercept, Figure 6a). Similarly, Group_-R + CF_ exhibited higher intercepts for vel_max_ and vel_min_ (Group_-R + CF_, vel_max_ = 30.4803, CI = [29.6730 31.2876], Group_-NR + NF_, vel_max_ = 24.5439, CI = [23.8895 25.1983], Intercept, Figure 6b; Group_-R + CF_, vel_min_ = 6.4902, CI = [6.1319 6.8486], Group_-NR + NF_, vel_min_ = 3.7302, CI = [3.4472 4.0132], Intercept, Figure 6c). Additionally, we found that Group_-R + CF_ also showed higher levels of movement fusion (Group_-R + CF_, FI = 1.5378, CI = [1.4604 1.6151], Group_-NR + NF_, FI = 1.0930, CI = [1.0260 1.1599], Intercept, Figure 6d). Importantly, we observed that Group_-R + CF_ exhibited steeper learning curves across all measures on day 1 (Group_-R + CF_, MT = -0.0036, CI = [-0.0040 -0.0032], Group_-NR + NF_, MT = -0.0018, CI = [-0.0023 -0.0012], Slope, Figure 6a; (Group_-R + CF_, vel_max_ = 0.0230, CI = [0.0192 0.0268], Group_-NR + NF_, vel_max_ = 0.0045, CI = [0.0017 0.0074], Slope, Figure 6b; (Group_-R + CF_, vel_min_ = 0.0217, CI = [0.0192 0.0242], Group_-NR + NF_, vel_min_ = 0.0075, CI = [0.0058 0.0092], Slope, Figure 6c; (Group_-R + CF_, FI = 0.0034, CI = [0.0029 0.0038], Group_-NR + NF_, FI = 0.0013, CI = [0.0009 0.0016], Slope, Figure 6d). These results replicate our findings from experiment 1 and highlight that explicit reward in combination with performance-based feedback both invigorates performances during early training and enhances learning leading to additional performance gains across training. Furthermore, results from experiment 2 showed that performance can be further improved across an additional testing day only if explicit reward in combination with performance-based feedback was provided (Group_-R + CF_, MT = -0.0020, CI = [-0.0022 -0.0017], Group_-NR + NF_, MT = -0.0011, CI = [-0.0015 -0.0007], Slope, Figure 6a; (Group_-R + CF_, vel_max_ = 0.0181, CI = [0.0150 0.0211], Group_-NR + NF_, vel_max_ = 0.0016, CI = [-0.0008 0.0041], Slope, Figure 6b; (Group_-R + CF_, vel_min_ = 0.0173, CI = [0.0153 0.0192], Group_-NR + NF_, vel_min_ = 0.0046, CI = [0.0030 0.0062], Slope, Figure 6c; (Group_-R + CF_, FI = 0.0020, CI = [0.0017 0.0022], Group_-NR + NF_, FI = 0.0008, CI = [0.0005 0.0011], Slope, Figure 6d).

**Figure 6.**
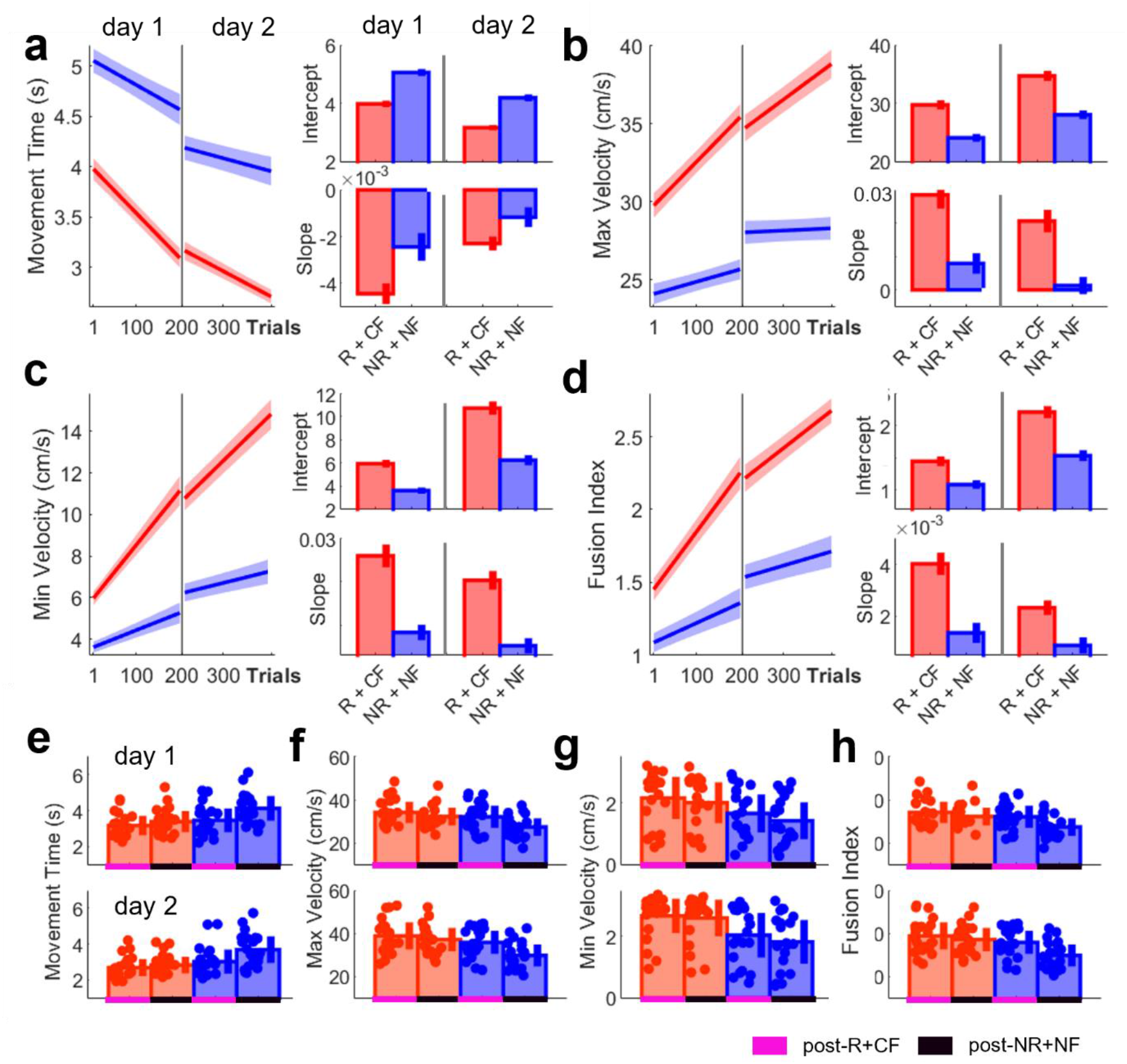
Explicit reward and correct performance feedback invigorates initial performance and enhances learning across both testing days. **a-d)** Averaged predicted model fits (simple polynomial model) to bootstrap estimates for each group including Intercept and Slope values (error bars represent 95% confidence intervals) for **a)** MT **b)** vel_max_ **c)** vel_min_ and **d)** FI. **e-h)** Post assessment performance comparing group performance during post-R+CF and post-NR+NF during day 1 and day 2 (upper and lower panel respectively) for **e)** MT **f)** vel_max_ **g)** vel_min_ and **h)** FI (shaded regions/error bars represent SEM).

Across post assessments, we found significant interactions between timepoint (post-R+CF vs post-NR+NF) and group (Group_-R + CF_ vs Group_-NR + NF_) on both days for MT (mixed ANOVA; day 1: F = 18.07, p < 0.0001; day 2: F = 19.99, p < 0.0001) and vel_max_ (mixed ANOVA; day 1: F = 8.02, p = 0.0072; day 2: F = 24.92, p < 0.0001). Specifically, there was a significant group difference during post-NR+NF for both MT (Wilcoxon test; day 1: Z = -2.82, p = 0.0192; day 2: Z = -3.27, p = 0.0044; Figure 6e) and vel_max_ (Wilcoxon test; day 1: Z = 2.84, p = 0.018; day 2: Z = 3.07, p = 0.0084; Figure 6f). However, during post-R+CF no differences were found (MT, Wilcoxon test; day 1: Z = -1.13, p = 1; day 2: Z = -1.38, p = 1; vel_max_, Wilcoxon test; day 1: Z = 0.86, p = 1; day 2: Z = 0.93, p = 1). This indicates that Group_-NR + NF_ were able to instantaneously invigorate their performance during post-R+CF. However, these performance gains were not maintained during post-NR+NF, suggesting that they remained transient in nature. Additionally, we found a significant effect for group on both testing days for vel_min_ (mixed ANOVA; day 1: F = 4.90, p = 0.0327; day 2: F = 7.70, p = 0.0083, Figure 6g) and FI (mixed ANOVA; day 1: F = 4.91, p = 0.0324; day 2: F = 7.38, p = 0.0097, Figure 6h). This suggests that the improvements in vel_min_ and FI within Group_-R + CF_ were more stable across post assessments (i.e. when reward was no longer available).

### Spatial reorganisation identifies the final stages of movement fusion and can be enhanced through explicit reward in combination with performance-based feedback

The results from both experiments suggest that movement fusion represents a viable strategy to further enhance performance (via increases in reaching speed (vel_max_) and decreases in dwell time in between reaches (vel_min_)). Importantly, movement fusion is a training-dependent process which can be accelerated by providing both explicit reward and performance-based feedback. Additionally, our findings indicate that fusion is not only associated with performance gains (i.e., faster MTs) during training, but also during periods without feedback (post-NR+NF). This may indicate that movement fusion allows for improved retention. To assess whether movement fusion led to a change in how the action is performed spatially, we assessed the radial distance (RD) between vel_max_ on the sub-movement and vel_min_ around the via points (Figure 7a). When executing a fluid point-to-point reaching movement (i.e., stopping in the target) vel_max_ will be spatially located approximately halfway through the reaching movement (Figure 7a right panel). However, with movement fusion vel_max_ will drift closer to vel_min_ which is located close to the target (Figure 7a left panel). Therefore, RD between the two becomes smaller with increasing FI levels, which can be expressed as a percentage of distance covered (Supplementary Figure 3). Hence, higher RD values represents the two movements merging together. We found that across training Group_-R+CF_ expressed spatial reorganisation as seen in a significant decrease in RD between vel_max_ and vel_min_ (Figure 8b).

**Figure 7.**
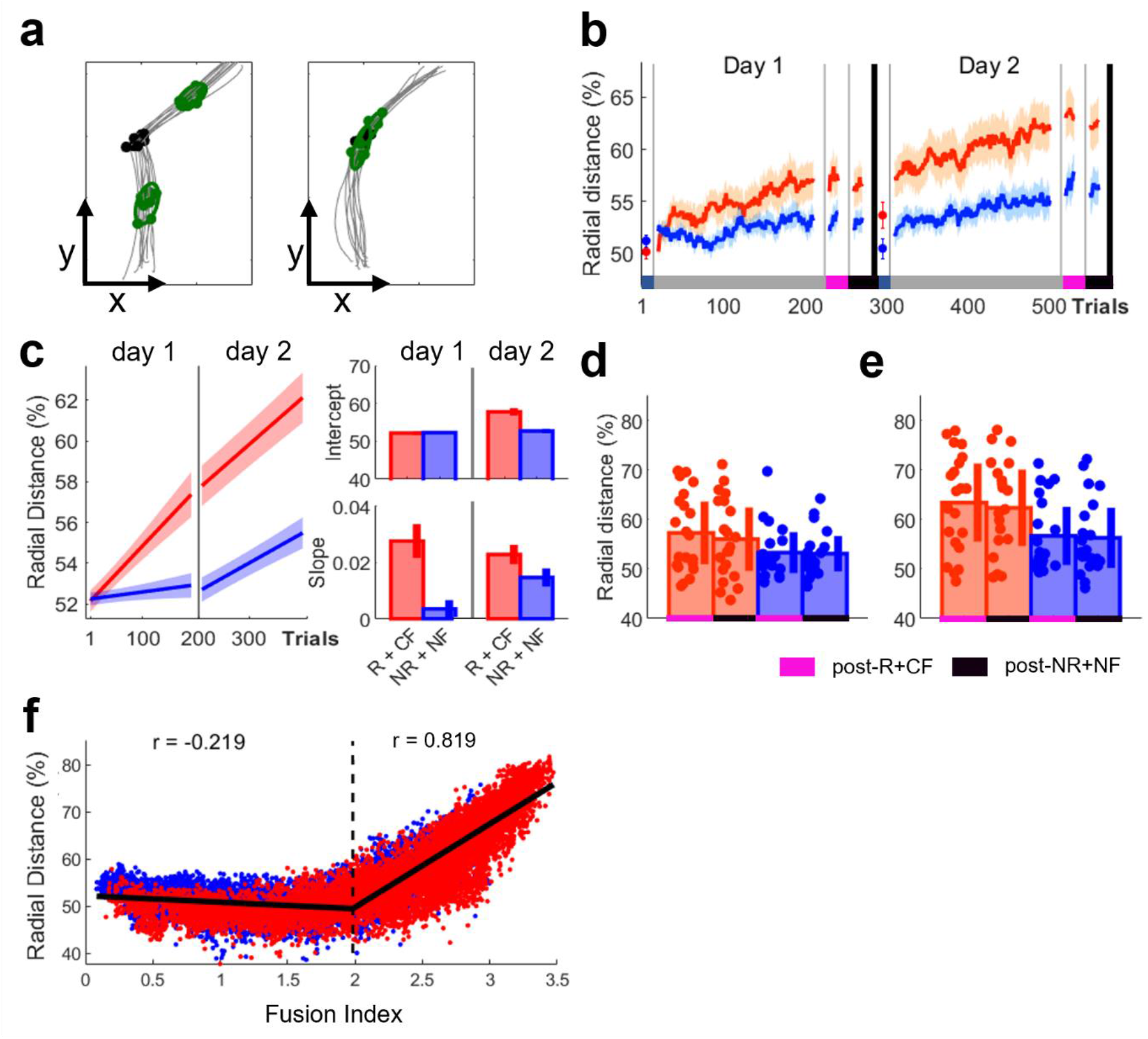
Reward-based improvements in spatial reorganisation. **a)** Example data for the spatial location of v_max_ when performing two individual (left panel) and one fused (right panel) reaching movement. **b)** Trial-by-trial changes in radial distance averaged over participants for both groups. **c)** Averaged predicted model fits (simple polynomial model) to bootstrap estimates for each group including Intercept and Slope values (error bars represent 95% confidence intervals). **d-e)** Post assessment comparing group performance during post-R+CF and post-NR+NF during **d)** Day 1 and **e)** Day 2 (shaded regions/error bars represent SEM). **f)** Scatterplot illustrating the relationship between mean FI levels and spatial reorganisation (radial distance %). It includes a two-segment piecewise linear function fitted to the data.

**Figure 8.**
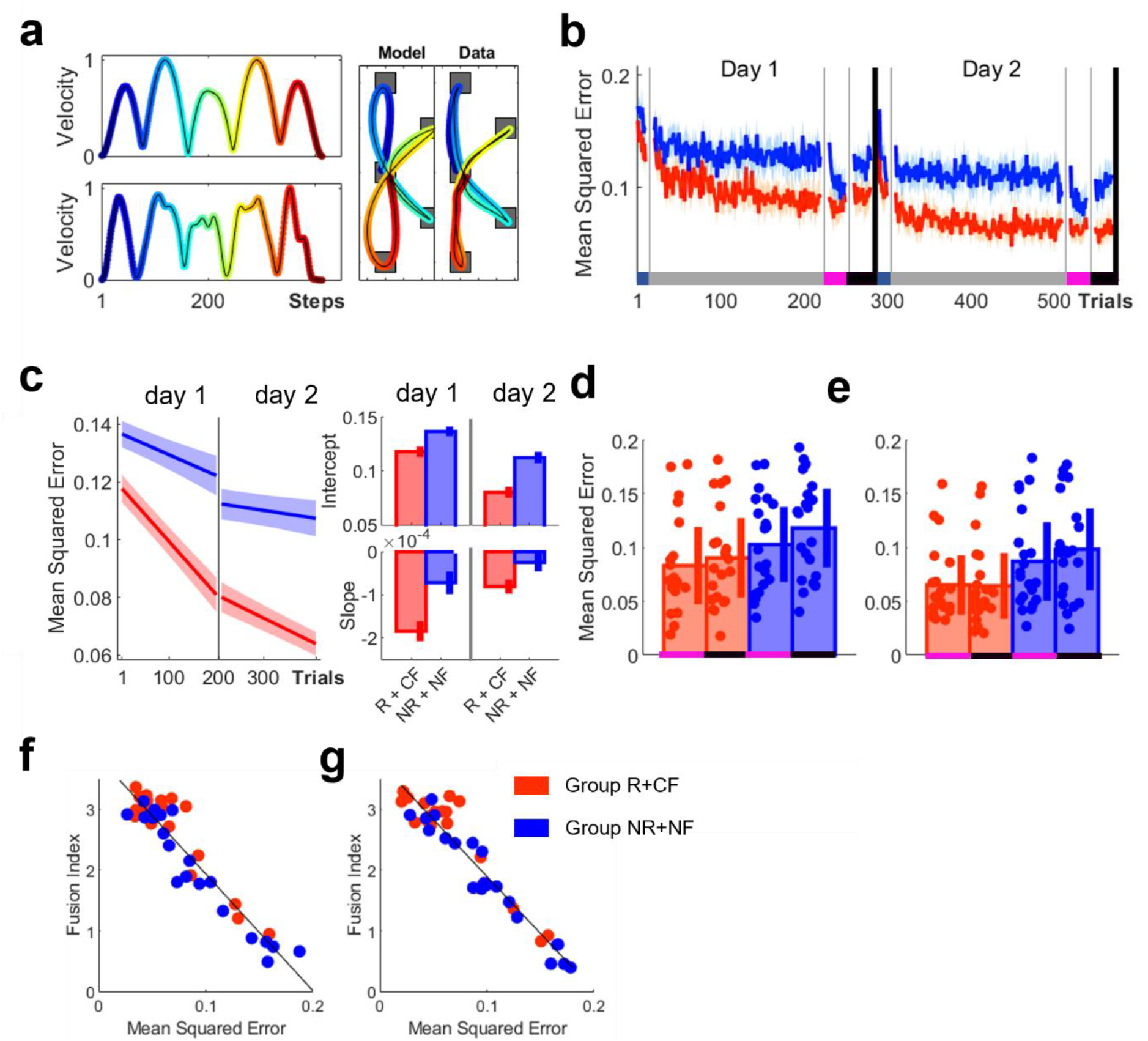
Movement fusion is associated with improvements in movement smoothness. **a)** Comparisons between data and the predictions of a minimum jerk model for both trajectory (right panel) and velocity profiles (left panel) for a single trial. **b)** Trial-by-trial changes in mean square error averaged over participants for both groups. **c)** Averaged predicted model fits (simple polynomial model) to bootstrap estimates for each group including Intercept and Slope values (error bars represent 95% confidence intervals). **d-e)** Post assessment comparing group performance during post-R+CF and post-NR+NF during **d)** Day 1 and **e)** Day 2 (shaded regions/error bars represent SEM). **f)** Scatterplot illustrating the relationship between mean FI levels and MSE for **f)** post-R+CF and **g)** post-NR+NF (pool across days).

No difference between groups in RD was observed during baseline (Wilcoxon test; Z = -1.18, p = 0.2371, Figure 7b). Instead, we found that explicit reward in combination with performance-based feedback enhanced performance during early training only on day 2 (day 1; Group_-R + CF_, RD = 52.0447, CI = [51.5142 52.5752]; Group_-NR + NF_, RD = 52.2428, CI = [51.9106 52.5751]; day 2; Group_-R + CF_, RD = 57.8621, CI = [56.8473 58.8768]; Group_-NR + NF_, RD = 52.7398, CI = [52.1450 53.3346]; Intercept, Figure 7c). This suggests spatial reorganisation is a learning-dependent process that cannot be instantly enhanced. Importantly, we found that Group_-R + CF_ exhibited a pronounced learning-related increase in RD across both testing days 2 (day 1; Group_-R + CF_, RD = 0.0277, CI = [0.0215 0.338]; Group_-NR + NF_, RD = 0.0036, CI = [0.0005 0.0067]; day 2; Group_-R + CF_, RD = 0.0232, CI = [0.0198 0.0267]; Group_-NR + NF_, RD = 0.0144, CI = [0.0112 0.0175]; Slope, Figure 7c).

Across post assessments, we found a significant main effect for group on day 2 (mixed ANOVA; day 1: F = 3.08, p = 0.0868; day 2: F = 5.76, p = 0.0211, Figure 6d, e). However, no main effect for timepoint was found (day 1: F = 1.87, p = 0.1796; day 2: F = 2.85, p = 0.0989). This suggest that changes in RDs were stable and more pronounced in Group_-R + CF_. To understand the relationship between FI and spatial reorganisation, we plotted them against each other and detected a pronounced drift in RD (%) with increasing FI levels resulting in a curvilinear shape (Figure 7f). After fitting a two-segment piecewise linear function to the data, we found an inflection point at ∼1.66 (FI) and a strong correlation between I and D for the second se ment (partial correlation controllin for roup; se ment 1: ρ = -0.16; p < 0.0001; segment 2: ρ = .89; p <. 1). his su ests that in order to fully fuse two consecutive movements, spatial reorganisation is required^26–28^ and this process can be enhanced with a combination of explicit reward and correct performance-based feedback.

### Movement fusion is associated with improvements in smoothness

As movement fusion involves the difference between vel_max_ and vel_min_ decreasing (Figure 5b), it implies that periods of acceleration/deceleration should become less pronounced and the movement ought to become smoother. To assess whether movement fusion is associated with increases in smoothness, participants’ performance was compared to the predictions of an optimisation model that minimised jerk across the movement sequence^30^. On trial-by-trial basis, mean squared error was calculated between the model and the actual velocity profile (Methods, Equation 4; Figure 8a). In summary, Group_-NR+NF_ became more aligned to the model’s predictions suggesting this roup’s performance became smoother (Figure 8b).

No difference between groups in mean squared error (MSE) was observed during baseline (Wilcoxon test; Z = -1.16, p = 0.2472). Instead, we found that explicit reward in combination with performance-based feedback enhanced performance during early training on both days (day 1; Group_-R + CF_, MSE = 0.1178, CI = [0.1130 0.1227]; Group_-NR + NF_, MSE = 0.1372, CI = [0.1326 0.1417]; day 2; Group_-R + CF_, MSE = 0.0796, CI = [0.0743 0.0850]; Group_-NR + NF_, MSE = 0.1129, CI = [0.1078 0.1180]; Intercept, Figure 8c). These results were in line with the movement fusion results and highlighted that R+CF enhanced movement smoothness during early training. Importantly, we found that Group_-R + CF_ exhibited a pronounced learning-related decrease in MSE across both testing days (day 1; Group_-R + CF_, MSE = -0.0018, CI = [-0.0021 -0.0016]; Group_-NR + NF_, MSE = -0.0007, CI = [-0.0010 -0.0005]; day 2; Group_-R + CF_, MSE = -0.0008, CI = [-0.0010 -0.0006]; Group_-NR + NF_, MSE = -0.0003, CI = [-0.0005 -0.0001]; Slope, Figure 8c). Additionally, during both post-R+CF (Figure 8f) and post-NR+NF (Figure 8g) we found that higher FI values are strongly associated with reduced MSE while accounting for the factor group (post-R+CF; rho = -0.944, p < 0.0001; post-NR+NF; rho = -0.952, p < 0.0001). This suggests that increases in movement fusion were related to improvements in smoothness. Further support comes from analysis which showed R+CF also reduced spectral arc length, an alternative measure of smoothness (See Methods^34,35^) (Supplementary Figure 4).

Across post assessments, we found a significant main effect for group on day 2 (mixed ANOVA; day 1: F = 3.08, p = 0.0868; day 2: F = 5.76, p = 0.0211, Figure 6d, e). However, no main effect for timepoint was found (day 1: F = 1.87, p = 0.1796; day 2: F = 2.85, p = 0.0989), which suggests that changes in RDs were stable and more pronounced in Group_-R + CF_.

### Performance gains are maintained across an additional testing day without reward

We next aimed to assess the robustness of these performance gains in an experiment including an additional testing day without any explicit reward and performance feedback (elongated washout condition; N=5). Participants underwent the same regime as in experiment 2 and on day 3 were asked to complete 200 no reward/feedback trials. Even after 24 hours, and over the course of 200 additional NR+NF trials, participants maintained similar MT performance levels. We used a repeated measures ANOVA with timepoint (early (first 15 trials) vs late (last 15 trials) across all testing days) as the within factor to assess changes across testing days (repeated measures ANOVA, F = 28.65, p < 0.0001; Figure 9a). These results indicate that performance improved over the course of the first 2 days (i.e. when reward was provided). However, no changes in MT performance could be observed between late training on day 2 and early training on day 3 (Wilcoxon test, Z = -1.21, p = 0.3016) and between early and late training on day 3 (Wilcoxon test, Z = -1.48, p = 0.4444).

**Figure 9.**
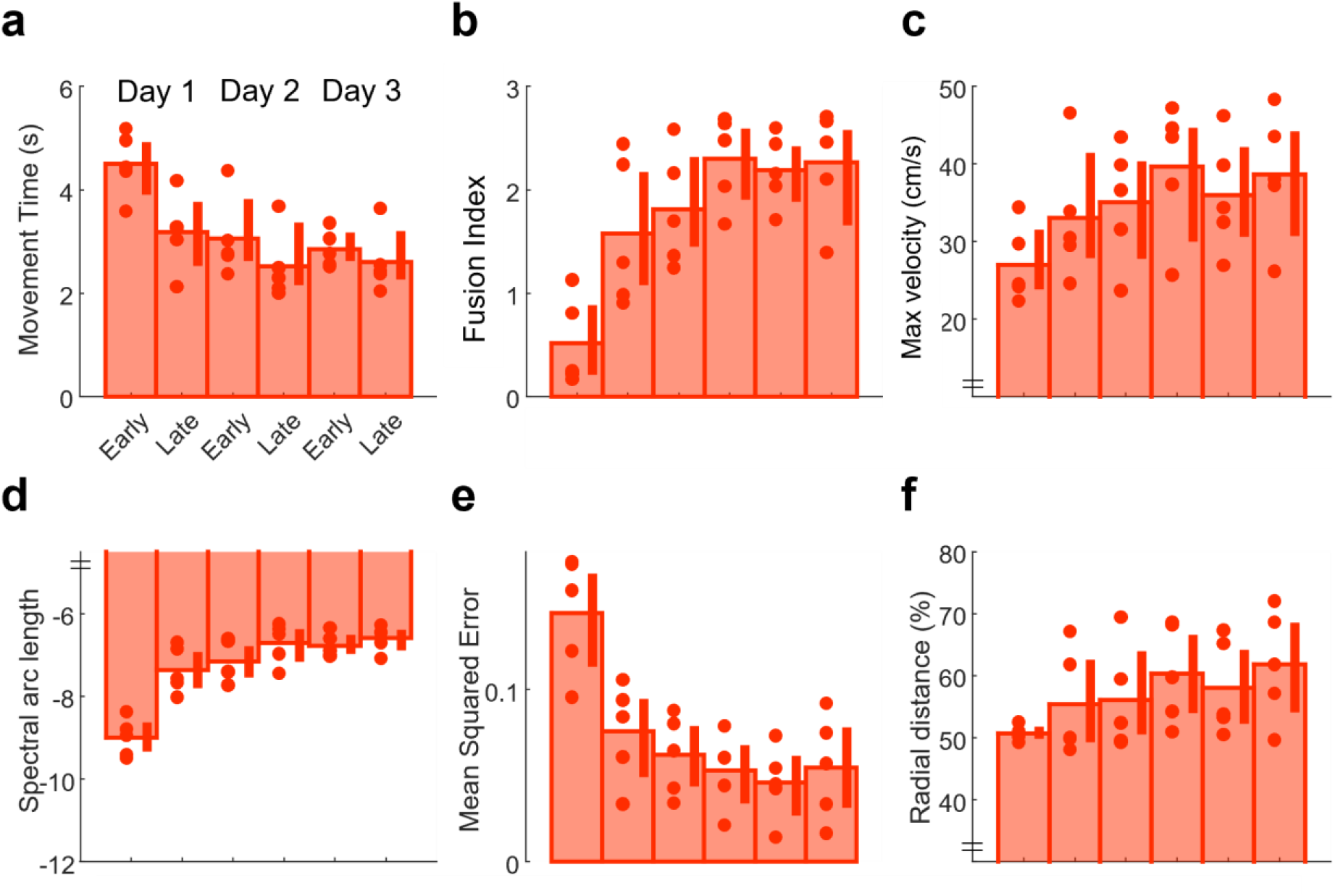
Long-term maintenance of performance without feedback. **a-f: Experiment 3**. Median data across all 3 days (early (first 20 trials) vs late (last 20 trials) training). **a)** MT **b)** FI level **c)** velmax **d)** Spectral arc length (smoothness) **e)** Mean squared error between data and minimum jerk model prediction **f)** Radial distance (spatial reorganisation). Shaded regions/error bars represent SEM. See Supplement Figure 5 for trial-by-trial data.

Similarly, fusion levels appeared stable across the additional testing day without feedback (repeated measures ANOVA, F = 19.19 p < 0.0001; Figure 9b), with no changes in performance between late day 2 and early day 3 (Wilcoxon test, Z = -0.94, p = 1) and no changes across day 3 (Wilcoxon test; early vs late; day 3, Z = -0.67, p = 1). In addition, vel_max_ were maintained transitioning to and across day 3 (Wilcoxon test, late training day 2 x early training day 3, Z = -1.21, p = 1; early training day 3 x late training day 3, Z = -1.21, p = 1; Figure 9c). When assessing changes in smoothness using spectral arc length, we found similar results to experiment 2. Smoothness improved over the course of the experiment (repeated measures ANOVA, F = 57.14 p < 0.0001; Figure 9), while no changes could be observed between late training on day 2 and early training on day 3 (Wilcoxon test, Z = -0.67, p = 1) and between early and late training on day 3 (Wilcoxon test, Z = -1.21, p = 1). In line with these results, we found that performance aligned progressively with the predictions of the minimum jerk model (repeated measures ANOVA, F = 23.38 p < 0.0001; Figure 9), while no significant changes in similarity could be observed transitioning to and across day 3 (Wilcoxon test, late training day 2 x early training day 3, Z = -2.02, p = 1; early training day 3 x late training day 3, Z = -1.21, p = 1). In addition, we found that participants progressively reorganised their spatial movement output, with a significant decrease in radial distance found across the experiment (repeated measures ANOVA, F = 57.14 p < 0.0001; Figure 9f). However, no changes in radial distance were observed between late training on day 2 and early training on day 3 (Wilcoxon test, Z = -2.02, p = 1) and between early and late training on day 3 (Wilcoxon test, Z = -1.75, p = 1). These findings supported our results from experiment 2 and may indicate that improvements in movement efficiency, through a quantitative change in how the task is performed, enables long-term retention of reward-based performance gains.

## Discussion

Previous research on the effects of reward on motor behaviour found that reward invigorated performance^1,3–5,11–14^ or enhanced motor learning and/or retention^8–10,15–17,36^. More specifically, studies using saccadic or discrete reaching movements have consistently shown that reward instantaneously improved MTs while maintaining similar levels of accuracy^1,3–5,11–14^, while most research employing force and button-press tasks found reward-related improvements in learning and/or retention (i.e., reductions in error rates or increases in number of successful trials)^8–10,15–17^. Our results suggest that these reward-related effects represent dissociable mechanisms and can be attributed to explicit reward and performance feedback, respectively. Experiment 1 showed that explicit reward (monetary incentive) instantly reduced MT during early training, while accurate performance feedback led to training-related improvements in MT irrespective of reward availability. Crucially, combining explicit reward with performance feedback resulted in both an instant reduction and a learning-related improvement in MT which maximised performance gains.

The instantaneous effect of explicit reward on motor behaviour has previously been explained by reward paying the energetic cost of enhanced performance^1,3,4^. For example, faster discrete reaching movements under reward conditions have been associated with increased arm stiffness^4^. Although an attractively simple strategy, increased stiffness comes with a marked escalation in metabolic costs^37^. Interestingly, these effects of reward are transient in that they are no longer observed once reward is removed^1,3,4^. Therefore, explicit reward seems to enable the transient use of energetically demanding (or even cognitively demanding ^1,14^) control mechanisms. Neurally this may relate to reward altering the trial-by-trial excitability of several motor regions (dorsal premotor cortex, primary motor cortex), irrespective of past reward history^38^. In humans, it has even been shown that trial-by-trial primary motor cortex excitability reflects the subjective value of reward and also mediates its incentivized effects on motor performance^39^

In contrast, performance-based feedback led to training-dependent improvements in MT. These findings are in line with previous work using force and button-press tasks that showed learning-dependent improvements in performance^8–10,15–17^. Performance feedback, provided by either points or binary feedback, represents a reinforcement-based teaching signal (implicit reward signal)^18,19,21^ which provides information on how well a motor task has been completed (knowledge of performance) and has been shown to enhance other forms of motor learning^11,20–24^ and retention^9–11^. Interestingly, recent research has shown that subpopulations of neurons in the primary motor cortex develop over training which signal outcome information of a single trial but are independent of reward and movement kinematics^40^. This suggests that feedback information is present in motor cortices alongside neurons that encode reward, and could indicate that explicit reward and performance feedback have dissociable impact on the motor system.

Crucially, explicit reward in combination with accurate performance feedback resulted in both an instant reduction and a learning-related improvement in MT which maximised performance gains. These findings are in line with recent research showing that such a combination led to enhanced learning and improvements in retention in a pinch force reproduction task ^15^. It has been suggested that explicit reward augments exploitative behaviour after successful feedback (i.e., reproduce successful behaviour) and increases explorative behaviour after unsuccessful feedback (i.e., magnitude of behavioural change)^41^. In other words, explicit reward boosts the reinforcement learning occurring with performance feedback^15,41^. Interestingly, explicit reward in combination with random performance feedback did not lead to learning-related improvements. Within the context of reinforcement learning, an agent is believed to find better solutions by updating actions based on the feedback received^18^. Specifically, if an action yields more reward than expected, its value will increase. This learning process increases the likelihood of maximising future rewards and is mediated by the exploration-exploitation trade-off^42,43^. Our results suggest that this learning process depends on accurate credit assignment, with random performance feedback clearly impairing this learning process.

Combining explicit reward with correct performance feedback not only maximised performance gains but also led to a greater amount of movement fusion. Fusion describes the process of blending together a series of distinct movements into a single continuous action. However, previous work using a simpler sequential reaching task showed that fusion takes up to 8 days (1200-2000 trials)^26,27,44^. This highlights that movement fusion is characterised by a very slow learning process which is not simply the logical consequence of training. In contrast, participants in the present study showed clear fusion after a single training session (200 training trials) with this being maximised in Group_-R+CF_. Movement fusion was not only strongly associated with improvements in MT (via both increases in vel_max_ and reductions in dwell times) but also with increases in movement smoothness. Improvements in jerk/smoothness have been shown to reduce metabolic costs, thereby enhancing overall movement efficiency^31,32^. Experiment 2 showed that performance in Group_-R+CF_ was smoother and exhibited greater similarity to a minimum jerk trajectory through movement fusion. This suggests that movement fusion represents an effective strategy to perform faster, smoother and more energetically efficient movements^26–28,31,32^. Furthermore, experiment 2 and 3 demonstrated that performance gains in Group_-R+CF_ were maintained across post-NR+NF and an additional testing day without either explicit reward or performance feedback available. In line with recent work on arm stiffness during discrete reaching movements^4^, a reduction in metabolic costs via movement fusion may enable faster movements even when no explicit reward or performance feedback is available. Alternatively, the long-term retention of performance gains may also be explained within the context of associative learning^45,46^. According to this framework, repetitive pairing of fast MTs with explicit reward during training may induce an implicit association between two events that can remain even when reward was removed^45,46^. This in turn could account for the long-term retention of performance gains across an additional testing day.

Our results showed that movement fusion was associated with smoother and, with regards to minimising jerk, more efficient execution. Interestingly, reaching movements performed by stroke patients exhibit reduced smoothness ^47–50^, with increases in jerk being due to a decomposition of movement into a series of sub-movements ^47–50^. However, over the course of the recovery process, performance becomes smoother as these sub-movements are progressively blended ^47–50^. Considering this theoretical proximity to the concept of movement fusion, we speculate that stroke recovery and movement fusion may follow similar principles. Consequently, fusion facilitated by explicit reward in combination with performance feedback could be a powerful tool in stroke rehabilitation to promote smooth and efficient sequential actions which form an essential component of everyday life activities.

In summary, explicit reward and performance feedback have dissociable effects on motor behaviour. Importantly, pairing both maximised performance gains and accelerated the slow optimisation process of movement fusion which leads to stable improvements in the speed and efficiency of sequential actions.

## Methods

### Participants

121 participants (24 males; age range 18 - 35) were recruited to participate in three experiments, which had been approved by the local research ethics committee of the University of Birmingham. All participants were novices to the task paradigm and were free of motor, visual and cognitive impairment. Most participants were self-reportedly right-handed (N = 9 left-handed participants) and gave written informed consent prior to the start of the experiment. For their participation, participants were remunerated with either course credits or money (£7.5/hour) and were able to earn additional money during the task depending on their performance. Depending on the experiment, participants were pseudo-randomly allocated to one of the available groups.

### Experimental apparatus

All experiments were performed using a Polhemus 3SPACE Fastrak tracking device (Colchester, Vermont U.S.A; with a sampling rate of 110Hz). Participants were seated in front of the experimental apparatus which included a table, a horizontally placed mirror 25cm above the table and a screen (Figure 1a). A low-latency Apple Cinema screen was placed 25cm above the mirror and displayed the workspace and participants’ hand position (represented by a reen cursor – diameter 1cm). On the table, participants were asked to perform 2-D reaching movements. Looking into the mirror, they were able to see the representation of their hand position reflected from the screen above. This setup effectively blocked their hand from sight. The experiment was run using MATLAB (The Mathworks, Natwick, MA), with Psychophysics Toolbox 3.

### Task design

Participants were asked to hit a series of targets displayed on the screen (Figure 1b). Four circular (1cm diameter) tar ets were arran ed around a centre tar et (‘via tar et’). Startin in the via target, participants had to perform eight continuous reaching movements to complete a trial. Target 1 and 4 were displaced by 10cm on the *y-*axis, whereas Target 2 and 3 were 5cm away from the via target with an angle of 126 degrees between them (Figure 1b). To start each trial, participants had to pass their cursor though the preparation box (2×2cm) on the left side of the workspace, which triggered the appearance of the start box (2×2cm) in the centre of the screen. After moving the cursor into the start box, participants had to wait for 1.5s for the targets to appear. This ensured that participants were stationary before reaching for the first target. Target appearance served as the go-signal and the start box turned into the via target (circle). Upon reaching the last target (via target), all targets disappeared, and participants had to wait for 1.5s before being allowed to exit the start box to reach for the preparation box to initiate a new trial. Participants had to repeat a trial if they missed a target or performed the reaching order incorrectly. Similarly, exiting the start box too early either at the beginning or at the end of each trial resulted in a missed trial.

### Reward structure and feedback

In experiment 1, participants were randomly allocated to one of the five groups: (1) no reward and no performance feedback (Group_-NR+NF_), (2) no reward and accurate performance feedback (Group_-NR+CF_), (3) explicit reward and no performance feedback (Group_-R+NF_), (4) explicit reward and accurate performance feedback (Group_-R+CF_), and (5) explicit reward and random performance feedback (Group_-R+RF,_ Figure 1e). This design allowed us to systematically evaluate how explicit reward and performance feedback influence performance during a complex sequential reaching task.

Participants that received explicit reward were informed that faster MTs would earn them more money. Reward trials were cued using a visual stimulus prior to the start of the trial (Figure 1e). Once participants moved into the preparation box, the start box appeared in yellow (visual stimulus). In contrast, participants that were in a NR group were told to move as fast and accurately as possible and here the start box remained black. Performance feedback was provided after completing a trial while participants moved from the start box to the preparation box to initiate a new trial. Feedback was displayed on the top of the screen (i.e., ‘ p out of p’). We used a closed-loop design to calculate the feedback in each trial. To calculate this, we included the MT values of the last 20 trials and organised them from fastest to slowest to determine the rank of the current trial within the given array. A rank in the top three (<= 90%) returned a value of 5p, ranks >= 80% and <90% were valued at 4p; ranks >=60% and <80% were awarded 3p; ranks >=40% and < 60% earned 2p while 1p was awarded for ranks >=20% and < 40%. A rank in the bottom three (<20%) returned a value of 0p. When participants started a new experimental block, performance in the first trial was compared to the last 20 trials of the previously completed block. While participants in Group_-R+CF_ and Group_-R+RF_ were told that the performance feedback corresponds to money (i.e., 5p = 5 pence), Group_-NR+CF_ was informed that it refers to points (i.e., 5p = 5 points) that do not add money. In contrast to the CF groups Group_-R+RF_ received random feedback, which was not performance-based, but was drawn randomly from feedback given to participants in Group_-R+CF_. To this end, we strung together all reward values given to participants in this group and randomly chose a value for feedback in a given trial for Group_-R+RF_. Participants, therefore, received feedback which was similar in reward probability without corresponding to actual performance.

### Experiment 1 experimental procedure

In this experiment, we investigated whether explicit reward and performance feedback have dissociable effects on a sequential reaching task. The experiment included an initial learning phase prior to the start of the experiment as well as a baseline, training and two post assessments. Participants were pseudo-randomly allocated to either one of the five groups (N = 74, Figure 1c, d). *Learning*: We included a learning phase prior to the start of the experiment for participants to be able to memorise the reaching sequence. This allowed us to attribute any performance gains to improvements in execution rather than memory. Once participants waited 1.5s inside the start box, the targets appeared which were numbered clockwise from 1 to 4 starting with the central top target. Participants were also able to see a number sequence at the top left of the screen displaying the order of target reaches (1 – 3 – 2 - 4). Participants were instructed to hit the targets according to the number sequence while also hitting the via target in between target reaches. They had to repeat a trial if they missed a target or performed the reaching order incorrectly. Similarly, exiting the start box too early either at the beginning or at the end of each trial resulted in a missed trial. After a cued trial, participants were asked to complete a trial from memory without the number sequence or numbers inside the targets. If participants failed a no cue trial more than twice, cues appeared in the following trial as a reminder. After a maximum of 10 cue and 10 no cue trials participants completed this block. *Baseline*: Participants in both groups completed 10 baseline trials, which were used to assess whether there were any pre-training differences between groups. All roups were instructed to ‘move as fast and accurately as possible’, while no explicit reward or performance-based feedback was available. *Training:* Participants completed 200 training trials and received a combination of explicit reward and performance feedback depending on the group that they were assigned to. The explicit reward groups were informed that during this part they would be able to earn money depending on how fast they complete each trial (200 reward trials). *Post assessments*: After training, participants from all groups were asked to complete two post assessments (20 trials each); one with both explicit reward and accurate performance feedback available (post-R+CF) and one without either (post-NR+NF). The order was counter-balanced across participants.

### Experiment 2 experimental procedure

In this experiment (N=42), we aimed to partially replicate the results from experiment 1. In experiment 2, only Group-R + CF and Group-NR + NF were included to contrast the most beneficial feedback regime (i.e., monetary incentive with accurate performance feedback with its logical opposite). Additionally, we added a further testing day (day 2) to investigate whether movement fusion can be further enhanced with additional training. During day 2, participants underwent the same experimental protocol as day 1 (Figure 1c).

### Experiment 3 experimental procedure

In this experiment, we aimed to test how robust reward-driven performance gains were over an additional testing day without explicit reward and performance feedback. Participants (N = 5) underwent the same regime as the reward group in experiment 2 on the first two testing days. On the third testing day after baseline, participants were asked to complete 200 NR+NF trials.

### Data analysis

Analysis code is available on the Open Science Framework website, alongside the experimental datasets at: https://osf.io/62wcz/. The analyses were performed in Matlab (Mathworks, Natick, MA).

### Movement time (MT)

MT was measured as the time between exiting the start box and reaching the last target. This excludes reaction time, which describes the time between target appearance and when the participants’ start position exceeded 2cm In summary, explicit reward and performance feedback have dissociable effects on motor behaviour. Importantly, pairing both maximised performance gains and accelerated the slow optimisation process of movement fusion which leads to stable improvements in the speed and efficiency of sequential actions.

### Maximum and minimum velocity

Through the derivative of positional data (*x, y*), we obtained velocity profiles for each trial which were smoothed usin a aussian smoothin kernel (σ =2). he velocity profile was then divided into segments representing movements to each individual target (8 segments) by identifying when the positional data was within 2cm of a target. We measured the maximum velocity (*v*_*max*_) of each segment by finding the maximum velocity:

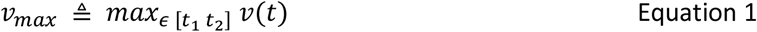

Where *v(t)* is the velocity of segment *t*, and *t*_*1*_ *and t*_*2*_ represent the start and end of segment *t* respectively. Similarly, minimum velocities (*v*_*min*_) were determined by measuring the minimum velocity when participants were inside a target (7 targets) using:

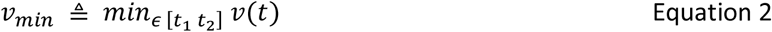

The individual maximum and minimum velocities were then averaged for each trial.

### Fusion index (FI)

Fusion describes the blending together of individual motor elements into a singular smooth action. This is represented in the velocity profile by the stop period between the two movements gradually disappearing and being replaced by a single velocity peak (Figure 4a, b) ^26–28^. To measure fusion, we compared the mean maximum velocities of two sequential reaches with the minimum velocity around the via point. The smaller the difference between these values, the greater coarticulation had occurred between the two movements (Figure 4b) ^51^. We calculated movement fusion by:

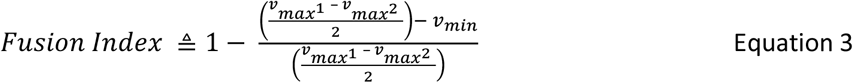

with *v*_*max*_^*1*^ and *v*_*max*_^*2*^ representing the velocity peak value of two reaching movements, respectively, and *v*_*min*_ representing the minimum value between these two points. We normalised the obtained difference, ranging from 0 to 1, with 1 indicating a fully coarticulated movement. Given that in this task seven transitions had to be completed, the maximum F value was 7 in each trial.

### Spatial reorganisation

In addition to FI, fusion can also be expressed spatially as the radial distance between the maximum velocity (v_max_) on the sub-movements and the minimum velocity (v_min_) around the via point (Figure 7a). This distance becomes smaller with increased movement fusion ^26,28^ and reflects the merging of two sub-movements into one (Supplementary Figure 3). To measure these changes in radial distance between v_max_ and v_min_, we used a sliding window approach of 10 trials at a time. For each target reach (excluding the first and the last) we fitted a confidence ellipse ^52^ with a 95% confidence criterion around the scatter of the spatial position (*x, y*) of each v_max_ of the included trials (Figure 7a). The confidence ellipses were obtained using principal component analysis to determine the minimum and maximum dispersion of the included data points in the *x-y* plane. To measure the distance between the scatter and its correspondin via point, we determined the ellipse’s centroid (point of intersection of ellipse’s axes) and calculated the radial distance to the via point. The obtained distance values were normalised and ranged from 0-100%, with 100% representing 0 cm distance between the centroid and the via point. Considering that individual reaching movements display a bell-shaped velocity profile, with the v_max_ situated approximately in the centre of the movement, radial distance values between 45-55% can be expected if each movement is executed individually (Supplementary Figure 3).

### Minimum-jerk model

A traditional minimum-jerk model for motor control is guided by optimisation theory, where a ‘cost’ is minimised over the trajectory ^29,30^. In the case of the minimum-jerk model, the cost is defined as the squared jerk (3^rd^ derivative of position with respect to time):

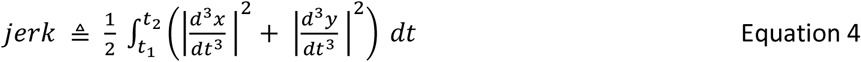

Here *x* and *y* represent the position of the index finger over time (*t*), while *t*_*1*_ and *t*_*2*_ define the start and end of a trial in seconds (*t*). The Matlab code provided by Todorov and Jordan (1998)^53^ was used to compute the minimum jerk trajectory (trajectory that minimised Equation 4), and the accompanying velocity profile, given a set of via points, start/end position and movement time ^30^. We then calculated the mean square error (immse function in Matlab) between the predicted and actual velocity profile, which were both normalised and interpolated (N = 500), to estimate the fit on a trial-by-trial basis. Due to the two-dimensional structure of trajectories, we used velocity profiles rather than the trajectories for this comparison.

### Spectral arc length

To further assess movement smoothness, we measured spectral arc length (see Supplementary Figure 4). Although we decided to use the traditional jerk metric in our modelling analysis, to allow for comparisons with prior literature, spectral arc length has been shown to be less sensitive to differences in MT and more sensitive to changes in smoothness ^35,54^. The spectral arc length is derived from the arc length of the power spectrum of a Fourier transformation of the velocity profile. We used an open-source Matlab toolbox to calculate this value for each trajectory ^55^.

For both spectral arc length and the minimum-jerk model, we only included non-corrected trials. Trials that were classified as corrected included at least one corrective movement to hit a previously missed target. These additional movements added peaks to the velocity profile which complicated model comparison and increased jerkiness disproportionally. Therefore, 1820 trials were excluded for both analyses (8.68% of all trials).

### Statistical analysis

ANOVAs (experiment 1) and Wilcoxon tests (experiment 2) were used to analyse differences in performance during baseline. To assess whether explicit reward and performance-based feedback have distinct effects on performance during training in Experiment 1 and 2 we computed 1000 bootstrap estimates of the data for each group. Each estimate represented a randomly generated data set (N=15, with N=14 for Group_-NR+CF_) with replacement. Specifically, one participant was randomly chosen from the group pool and added to the new data set. This participant was subsequently included into the group pool again before randomly selecting another participant to be added to the new data set. Therefore, the same participant could be added to the new data set multiple times. A simple polynomial model (*f(x) = p1*x + p2*) was then fit to the mean of the new trial-by-trial training data set (200 trials) for each of the 1000 bootstrap estimates of each group. The 95% confidence intervals for each model parameter were used to represent significant differences between groups^33^. While *p2* represents the performance intercept during early training (1^st^-15^th^ trial), *p1* corresponds to the gradient (learning rate) across training. This model and analysis was chosen as it provided a simple but powerful assessment of the dissociable instantaneous (intercept) and learning-related effects associated with the different forms of feedback across groups. Mixed model ANOVAs were used to assess statistical significance during the post assessments with condition (post-R+CF, post-NR+NF (all 20 trials in each)) and group (experiment 1: Group_-R+CF_, Group_-R+RF_, Group_-R+NF_, Group_-NR+CF_, and Group_-NR+NF_; experiment 2: Group_-R+CF_ and Group_-NR+NF_) as factors. Note, that the model analysis was not required here as performance was stable across trials. We used one-sample Kolmogorov-Smirnov tests to test our data for normality and found that all measures were non-parametric. Median values were therefore used as input in all mixed model ANOVAs (similar to^56^). Wilcoxon tests were employed when a significant interaction and/or main effects were reported and corrections for multiple comparisons were performed using Bonferroni correction. Linear partial correlations (fitlm function in Matlab) were used to measure the degree of association between the chosen variables, while accounting for the factor group. Piecewise linear spline functions were fitted through the scatter of spatial distance values and FI levels using least square optimisation by means of shape language modelling (SLM) ^57^. We used three knots as input for the linear model.

A repeated-measure ANOVA was used to test for significance of our results in experiment 3. We compared performance with timepoint (early training (first 15 trial), late training (last 15 trials) over all 3 testing days) as the within factor. Due to our data being non-parametric after using one-sample Kolmogorov-Smirnov tests, we included median values as input for all repeated-measure ANOVAs. Wilcoxon test was used as post-hoc test and multiple comparisons were corrected for using Bonferroni corrections. Repeated-measure ANOVAs were chosen to analyse performance in Experiment 3 because only Group_-R+CF_ was included which did not require a more complex group comparison.

## Acknowledgements

This work was supported by the European Research Council starting grant: MotMotLearn (637488)

## Supplement (5 figures)

**Supplementary Figure 1.**
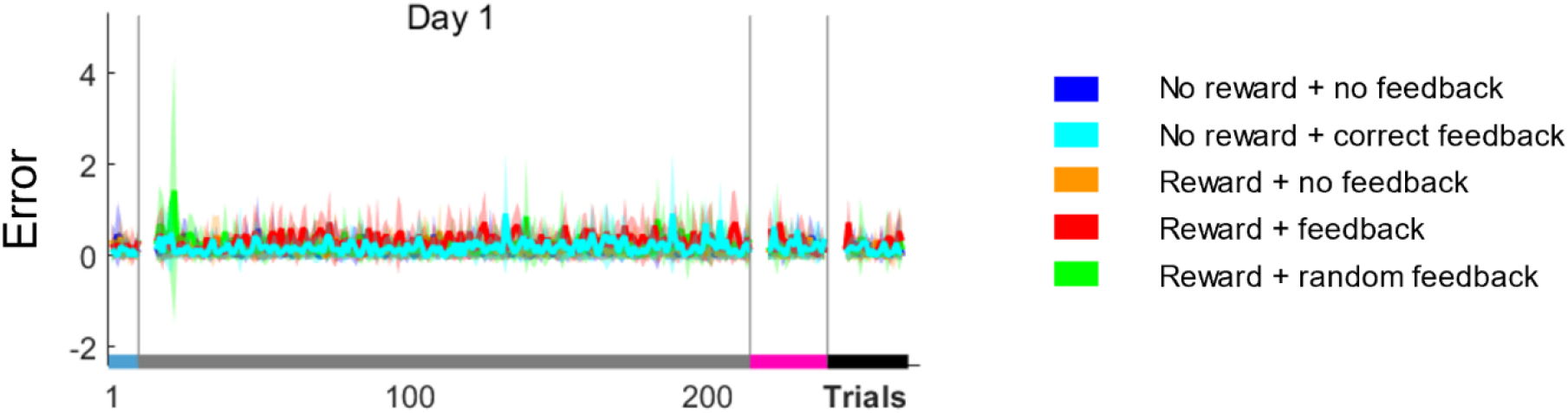
Trial-by-trial changes in performance averaged over participants for all groups in experiment 1. Averaged error rates across participants were similar across groups and were consistently below one error per trial.

**Supplementary Figure 2.**
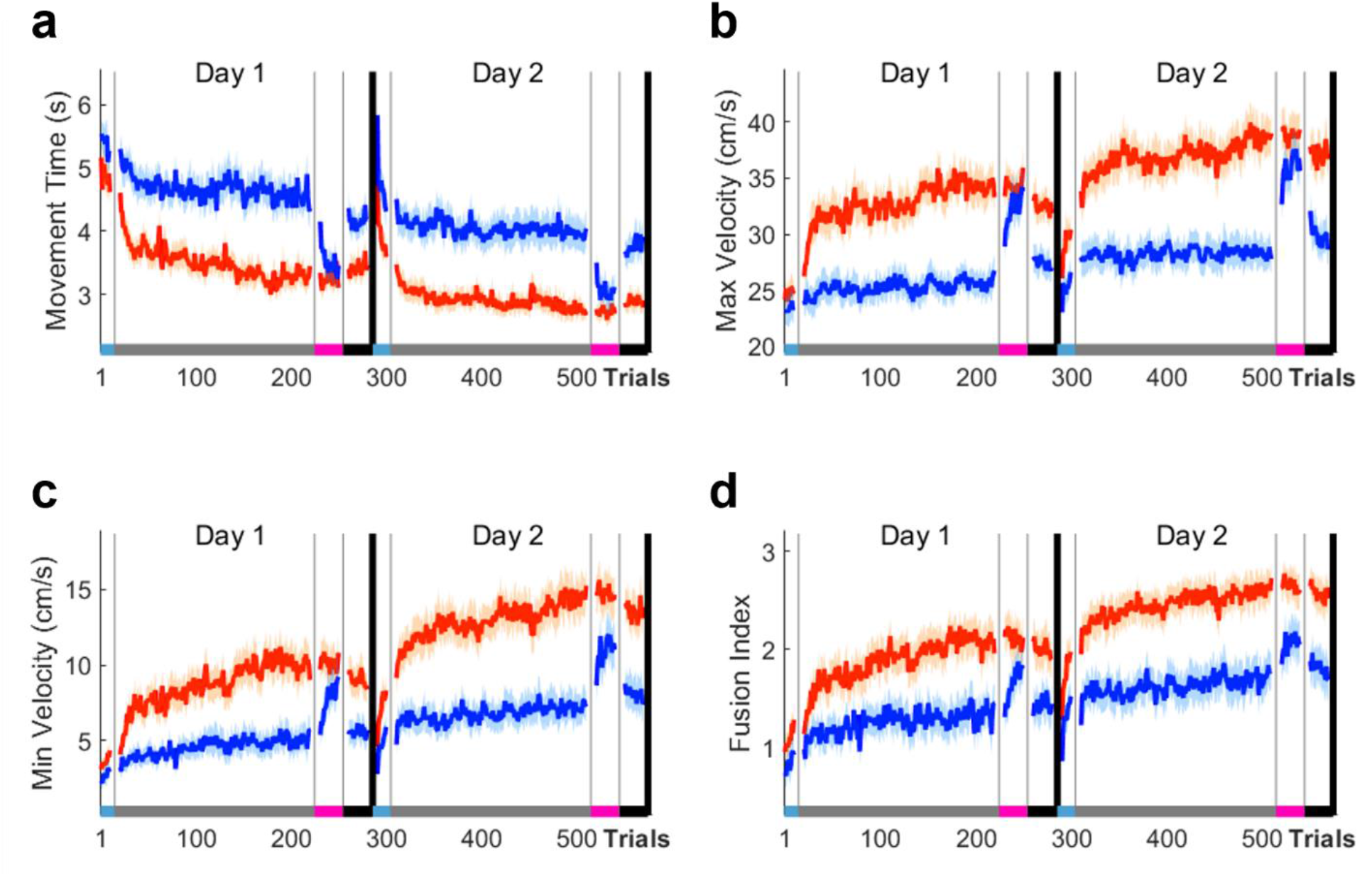
Trial-by-trial changes in performance averaged over participants for both Group_-R+CF_ nd Group_-NR+NF_. **a)** Movement time (MT), **b)** maximum velocity (vel_max_), **c)** minimum velocity (vel_min_) and **d)** fusion index (FI). Across outcome measures explicit reward in combination with correct performance feedback maximised performance gains.

**Supplementary Figure 3.**
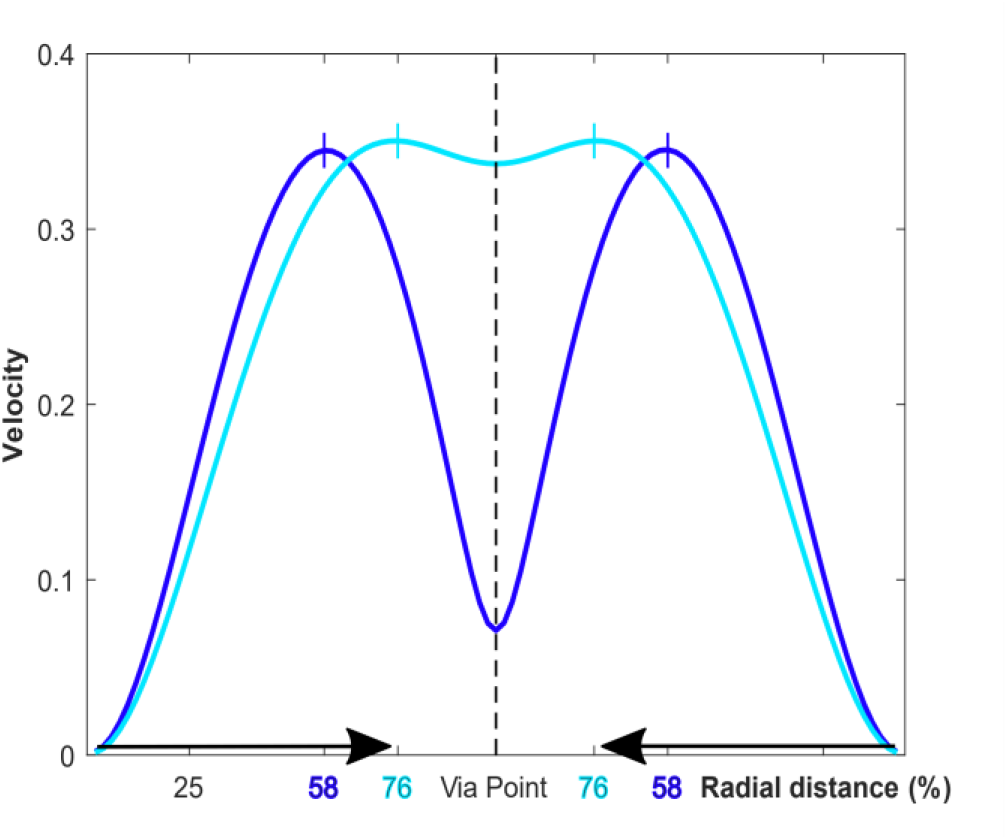
Illustration of radial distance (%) values depending on different levels of coarticulation. Considering two target reaches (same length) the spatial position (*x,y*) changes depending on the level of movement fusion thereby exhibiting less radial distance to the via point.

**Supplementary Figure 4.**
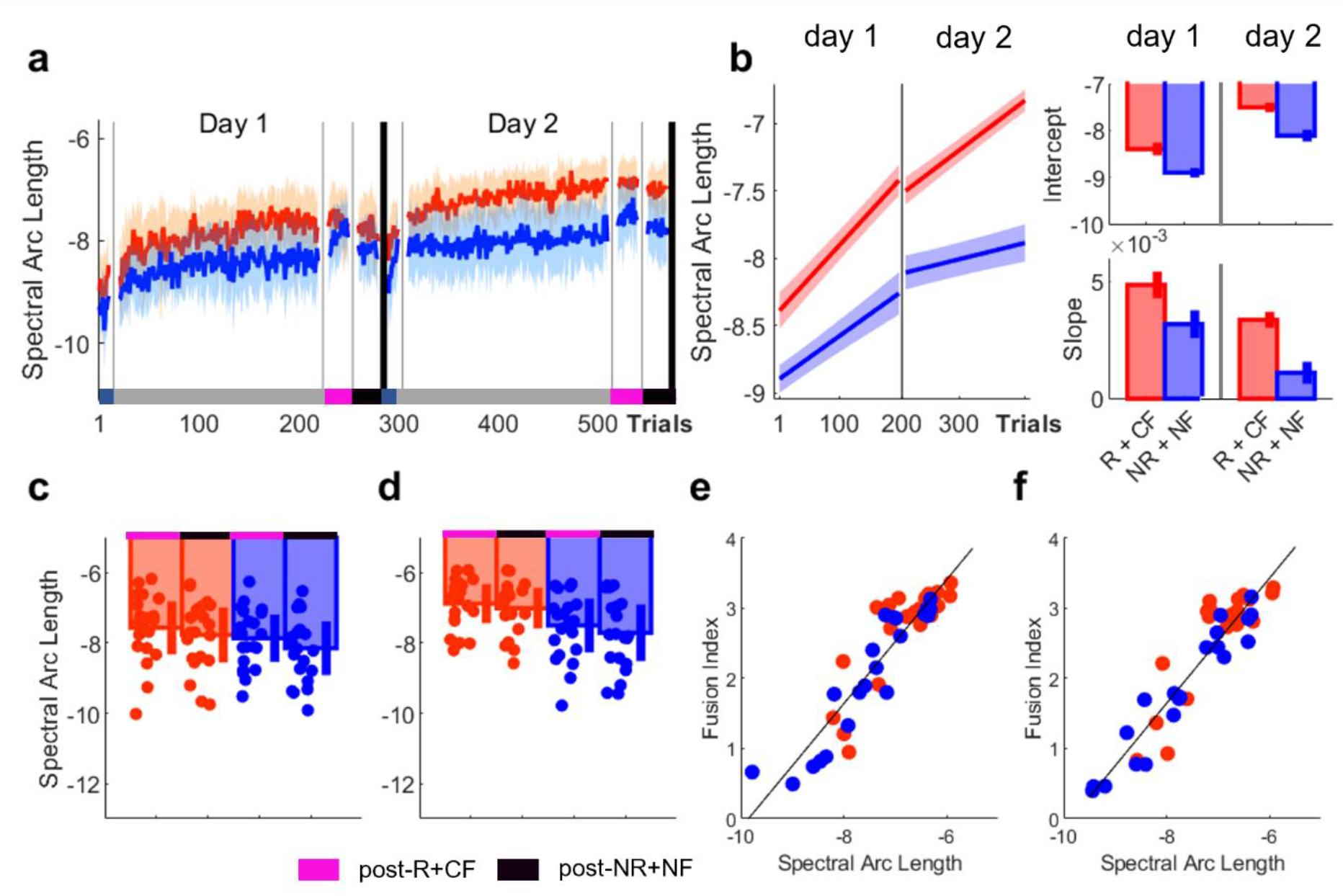
Movement fusion is strongly associated with improvements in movement smoothness. **a)** Trial-by-trial changes in spectral arc length averaged over participants for both groups. **b)** Averaged predicted model fits (simple polynomial model) to bootstrap estimates for each group including Intercept and Slope values (error bars and shaded region represent 95% confidence intervals). **c-d)** Post assessment comparing group performance during post-R+CF and post-NR+NF during **d)** Day 1 and **e)** Day 2 (error bars represent SEM). **f)** Scatterplot illustrating the relationship between mean spectral arc length and FI levels for **f)** post-R+CF and **g)** post-NR+NF (pool across days). No difference between groups in spectral arc length (SAP) was observed during baseline (Wilcoxon test; Z = 1.79, p = 0.0741, Supplementary Figure 3a). Instead, we found that explicit reward in combination with performance-based feedback enhanced performance during early training on both days (day 1; Group_-R + CF_, SAP = -8.3871, CI = [-8.5213 -8.2530]; Group_-NR + NF_, SAP = -8.8970, CI = [-8.9934 -8.8006]; day 2; Group_-R + CF_, SAP = -7.4975, CI = [-7.5977 -7.3973]; Group_-NR + NF_, SAP = -8.1021, CI = [-8.2277 -7.9766]; Intercept, Supplementary Figure 3b). These results were in line with both the movement fusion and model comparison results and highlighted that R+CF enhanced movement smoothness during early training. Importantly, we found that Group_-R + CF_ exhibited a pronounced learning-related decrease in SAP across both testing days (day 1; Group_-R + CF_, SAP = 0.0049, CI = [0.0043 0.0054]; Group_-NR + NF_, SAP = 0.0031, CI = [0.0025 0.0037]; day 2; Group_-R + CF_, SAP = 0.0034, CI = [0.0025 0.0037]; Group_-NR + NF_, SAP = 0.0010, CI = [0.0005 0.0015]; Slope, Supplementary Figure 3b). Across post assessments, we found a significant main effect for group on day 2 (mixed ANOVA; day 1: F = 1.44, p = 0.2369; day 2: F = 6.41, p = 0.0154, Figure 6d, e). Additionally, during both post-R+CF and post-NR+NF (Supplementary Figure 3e, f) we found that higher FI values are strongly associated with reduced SAP while accounting for the factor group (post-R+CF; rho = 0.897, p < 0.0001; post-NR+NF; rho = 0.921, p < 0.0001, Supplementary Figure 3e, f). In summary, using spectral arc length as a smoothness metric replicated the results from the minimum jerk model comparison.

**Supplementary Figure 5.**
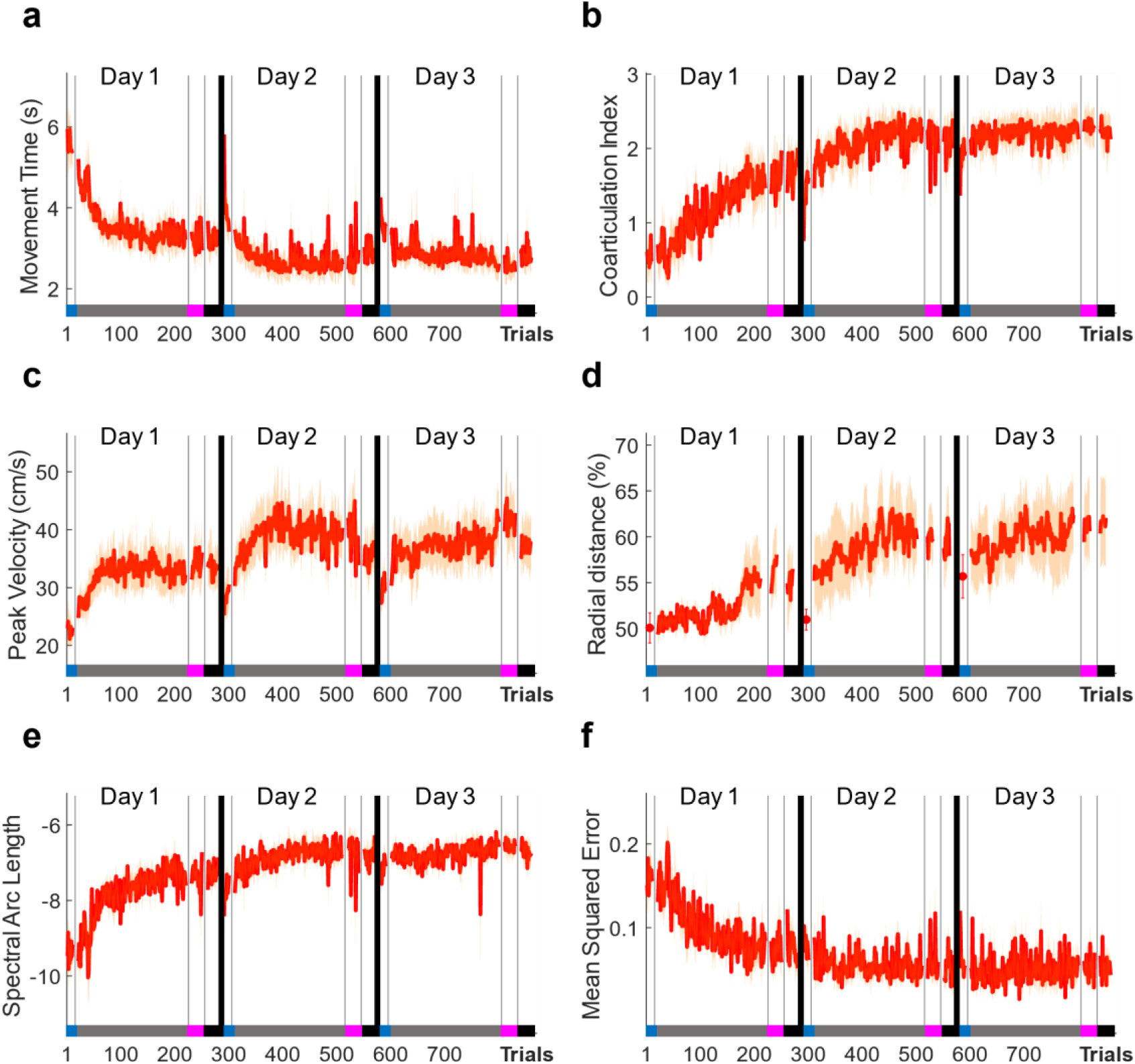
Performance gains are stable even over a prolonged washout period. Trial-by-trial data for **a)** MT, **b)** FI levels **c)** maximum velocity **d)** radial distance, **e)** spectral arc length, **f)** mean squared error for the third experiment which was scheduled on three consecutive days, with NR+NF scheduled for day 3.

